# The stress-induced transcription factor ATF4 has multiple conserved retrocopies that can alter gene expression

**DOI:** 10.64898/2026.03.31.714484

**Authors:** Hans M. Dalton, Elizabeth M Brydon, Tiffany S. Chan, Katie Owings, Mandi Wild, Naomi J. Young, Clement Y. Chow

**Affiliations:** Department of Human Genetics, University of Utah School of Medicine, Salt Lake City, Utah, United States of America; Department of Molecular Biosciences, University of Kansas, Lawrence, Kansas, United States of America

## Abstract

The cell must defend against various stressors from internal and external sources that disrupt cell homeostasis. The integrated stress response (ISR) is a highly conserved pathway that helps restore this homeostasis through upregulating the transcription factor, ATF4. Despite its importance to cell health and human disease, *ATF4* has several duplications in humans that have never been studied. Here, we characterize three retroduplications (retrocopies) of *ATF4* in humans for the first time. Evolutionary analysis demonstrates that these retrocopies are present and intact in many primate species over the past 37 million years, including several independent copies. We also find evidence of positive selection among primates. Human ATF4 retrocopies show basal transcription in healthy, unstressed cells and can be upregulated by the ISR. When translated in human cells, ATF4 retrocopy proteins are regulated by the proteasome in the same way as the parent ATF4 protein. Remarkably, each retrocopy can also alter the expression of several canonical ATF4 target genes, demonstrating that they can impact ISR-ATF4 stress signaling. Overall, *ATF4* retrocopies are conserved, biologically functional, and should be considered in future studies of the ISR and ATF4.

## Introduction

Cellular pathways like the integrated stress response (ISR) are activated in response to stress to restore homeostasis. The ISR is activated by viral infection, amino acid deprivation, endoplasmic reticulum stress, or heme deprivation, through the protein kinases PKR, GCN2, PERK, and HRI, respectively^1,2^. In response to this stress, the ISR inactivates the translation initiation factor eIF2B, downregulating global translation, while simultaneously upregulating specific translation of the ATF4 transcription factor^1,2^.

ATF4 is basic leucine zipper transcription factor that is critical to restoring cell homeostasis or, under high or prolonged stress, activating apoptosis^1,3,4^. It modulates the expression of hundreds of genes, including those involved in amino acid synthesis, autophagy, and apoptosis, among others^4–6^. ATF4 also has important roles in development^7–10^ and tumor biology^7,11–16^ in humans. The basic leucine zipper domain allows ATF4 to localize to the nucleus, bind to DNA, and dimerize with proteins such as CHOP and ATF3^17^.

ATF4 is regulated at the protein and mRNA level. ATF4 protein contains a p300 interaction domain that enables interaction with the histone acetyltransferase p300^18^. This interaction increases ATF4 stability and is independent of the p300 acetyltransferase activity^18^. ATF4 also contains two known degradation domains: an oxygen dependent degradation (ODD) domain and a ßTrCP motif^1^. Proline residues in the ODD domain are hydroxylated by the oxygen sensor PHD3, which are then targeted by the ubiquitin ligases Siah1 and Siah2^19,20^. The ßTrCP motif can be phosphorylated, leading to targeting by the SCF^βTrCP^ ubiquitin ligase^21–23^. Ultimately, each degradation domain leads to ubiquitination and degradation by the proteasome, and thus, ATF4 has a half-life of less than one hour^1,19,23^.

Under normal translation initiation, the small ribosomal subunit associates with the ternary complex (eIF2, GTP, and Met-tRNAi^Met^), binds to mRNAs, and scans for open reading frames (ORFs). Once an ORF is found, the ternary complex dissociates for the ribosome to form and initiate translation. The small ribosomal subunit must bind to a new ternary complex each time to re-initiate translation and build ribosomes. When the ISR is activated, eIF2B (part of the ternary complex) becomes inhibited and reduces available ternary complexes, inhibiting global protein synthesis. ATF4 uses transcript-level regulation to evade this inhibition by using three upstream open reading frames (uORFs)^24–26^. uORFs are nucleotide sequences, upstream or overlapping with the coding sequence (CDS) of a gene, that can affect its translation efficacy^27^. In *ATF4*, uORF1 is 6 nucleotides long, and uORF2 is 12 nucleotides long. uORF1 and 2 are located within the 5’ UTR, and do not overlap the CDS. In contrast, uORF3 is 180 nucleotides long and overlaps with the *ATF4* CDS. In normal, unstressed conditions, ribosomes first form on uORFs 1 or 2, and complete this short translation. Because ternary complexes are abundant under non-stressed conditions, the small ribosomal subunit continues scanning, and it usually finds and translates uORF3. Because uORF3 overlaps with the CDS, this process often “skips” over *ATF4* and consequently, ATF4 protein is rarely produced^24–26^. However, under the ISR, there are not enough ternary complexes to quickly re-initiate translation at uORF3, so scanning instead continues to the CDS ORF, at which point enough time has passed to allow a new ternary complex to bind. Thus, under the ISR, full length ATF4 is translated more frequently^24–26^. These uORFs are highly conserved in vertebrates, indicating their importance in ATF4 regulation^25^.

*ATF4* has four gene duplications in humans (annotated as pseudogenes *ATF4P1-P4*) that have never been characterized. While rare, gene duplications can maintain or even gain functionality^28–30^. One class is retroduplications (retrocopies), which are reverse transcribed (via transposon activity^31,32^) from the mRNA of “parent” genes and inserted back into the genome. When retrocopies are found to be functional, they are labeled as retrogenes^33^, such as retroCHMP3^34^, PTENP1^35,36^, and PGK2^37–39^. If a gene duplication is functional and beneficial, evolution can maintain them in the genome^28^. One major source of evolutionary pressure on the ISR comes from viruses^40,41^. For example, hepatitis B hijacks ISR signaling proteins in order to prevent downstream apoptosis^42^. Adenovirus produces decoy dsRNAs that sequester the ISR kinase PKR without activating it, thereby inhibiting its anti-viral activity^40,43^. These pressures have led to evolutionary selection in ISR machinery, such as in PKR, where it has evolved to evade viral mimicry^44^. Given that ATF4 is the critical downstream transcription factor of the ISR, such evolutionary pressure could select for these *ATF4* duplications if they influence ISR function.

Here, we demonstrate that the four *ATF4* duplications consist of three retrogenes and one duplicate retrogene. These retrogenes span evolutionary events across at least 37 million years in vertebrates. There is evidence of divergent and convergent evolution, with one *ATF4* retrogene truncation arising independently in at least three different primate species. Human retrogenes are expressed under normal conditions and ISR activation, are degraded similarly to *ATF4*, and can alter the expression of ATF4 target genes. Given their evolutionary history and function in cells, these retrogenes are likely important in the highly conserved ISR pathway in vertebrates.

## Results

### Humans have three retrocopies of *ATF4*

There are four annotated *ATF4* duplications: *ATF4P1, ATF4P2*, *ATF4P3*, and *ATF4P4*. All of these duplications lack introns and end in truncated PolyA tails - key characteristics of retrocopies^33,45^ (Fig. 1A, Supp Table 1). Retrocopies often originate from long interspersed element-1 (LINE-1) retrotransposons in mammals^31,32^, and these leave short target-site duplications (TSDs) flanking each gene^32,46^. We examined the region around each retrocopy and found evidence for TSDs in all of them (Supp Table 1). Thus, *ATF4P1-4* are retrocopies that emerged through LINE-1 retrotransposons.

**Figure 1.**
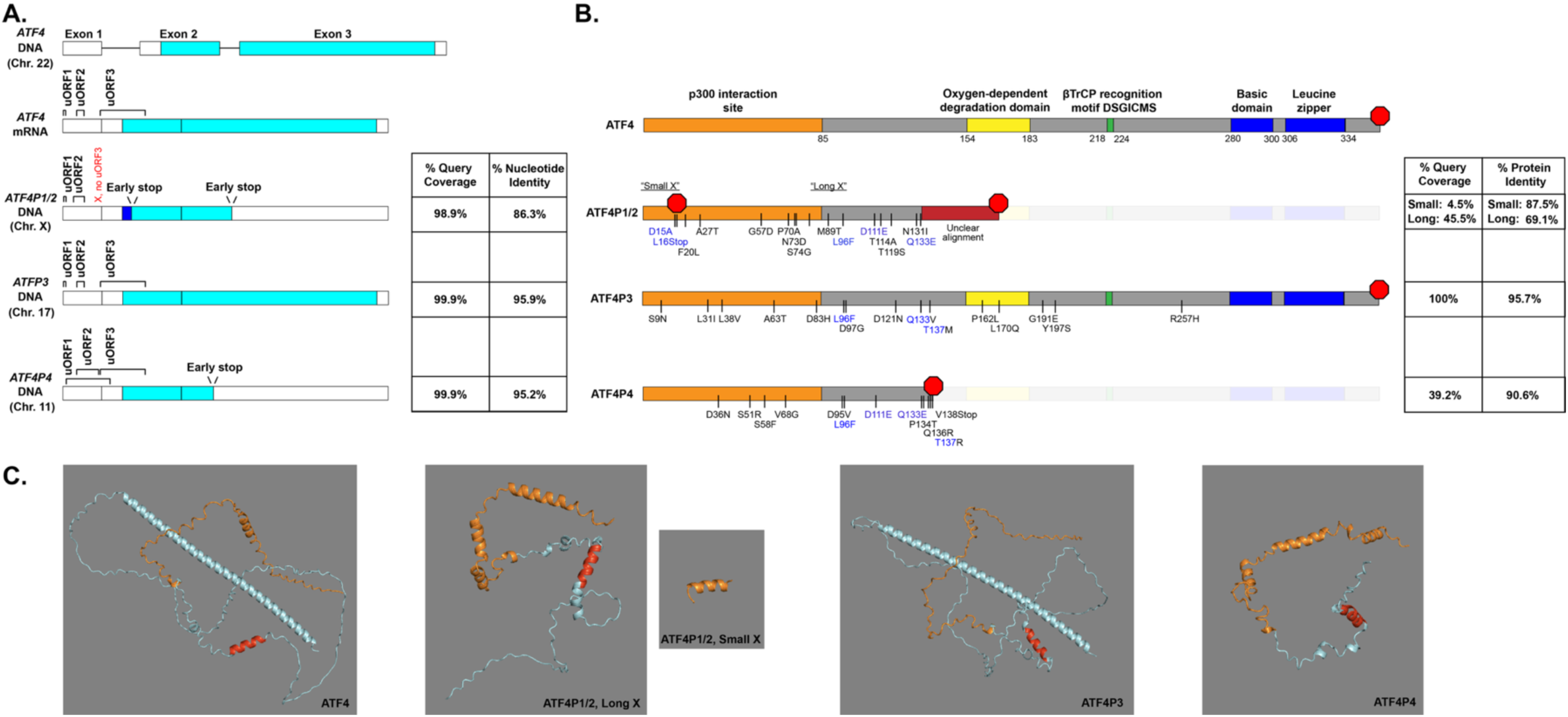
Characterizing four duplications of *ATF4*. **A.** Cartoons of genomic *ATF4*, *ATF4* mRNA, and genomic *ATF4P1-P4*. We list the current status of uORFs here, including when it is lost in *ATF4P1/P2*. We also list the % coverage and identity of each duplicate nucleotide sequence (minus flanking TSDs) when compared to *ATF4*. **B.** A diagram of each translated retrocopy compared to the translated Ensembl canonical transcript of *ATF4*. Amino acid changes that occur in more than one retrocopy are highlighted in blue. We also list the % coverage and identity of each translated protein sequence when compared to translation of *ATF4* mRNA (cDNA). **C.** Alphafold 3 predictions of translated retrocopy transcripts, visualized in PyMOL. Specific protein domains are colored for comparison: the p300 interaction domain (orange), an alpha helix structure that is maintained in all proteins except ATF4P1/2 Small X (red), and the rest of the protein including the basic leucine zipper domain (blue).

We compared the nucleotide sequence and putative protein translation of each retrocopy to their parent gene, *ATF4* (Fig. 1):

#### ATF4P2 and ATF4P1

*ATF4P2* and *ATF4P1* are located on the X Chromosome. Here, there is a larger ape-specific chromosomal reverse tandem duplication event^47^. The proximal border of this duplication is in the middle of the gene, *IKBKG*, resulting in the distal end containing a truncated duplication (the pseudogene, *IKBKGP1*). As *ATF4P1* is closer to the more recent *IKBKGP1*, it must be duplicated from *ATF4P2*, which is closer to *IKBKG*. This duplicated region is highly invariant. As such, human *ATF4P2* and *ATF4P1* are 100% identical at the nucleotide level from the first uORF to the PolyA tail (Supp Table 1), and we refer to them together as *ATF4P1/2* throughout.

*ATF4P1/2* maintain uORF1 and uORF2, but they have lost uORF3 due to a new stop codon (Fig. 1A). This very early premature stop codon would result in a putative 15 amino acid product. However, *ATF4* and *ATF4P1/2* have a secondary in-frame start site after 48 base pairs. This is immediately following the premature stop in *ATF4P1/2* and encodes a 170 amino acid product ending with multiple insertions which result in missense variants and a premature stop codon after the p300 interaction domain (Fig 1B-C). Going forward, we will refer to these products as “Small X” (15 amino acids) and “Long X” (170 amino acids), respectively. At 4.5% query coverage (15 amino acids), ATF4P1/2 Small X has 87.5% identity with ATF4. At 45.5% query coverage (170 amino acids), ATF4P1/2 Long X has 69.1% protein identity with ATF4 (Fig. 1B).

#### ATF4P3

*ATF4P3* is located on Chromosome 17 in an intron of the *RNF157* gene (Supp Fig 1). Because *ATF4P3* was inserted in the same strand orientation of *RNF157*, it might utilize the *RNF157* promoter or its regulatory sites. At the nucleotide level, all of its upstream open reading frames (uORFs) are intact (Fig. 1A), suggesting similar translational regulation to *ATF4*. *ATF4P3* encodes for a putative full-length ATF4 protein with zero nonsense changes. At 100% query coverage, ATF4P3 has 95.7% protein identity with ATF4 (Fig. 1B). Notably, the majority of its missense changes (12/15) occur in the first half of the gene, with fewer in the second half (3/15), and none are present in the basic leucine zipper domains (Fig. 1B). Because the ATF4 leucine zipper domain contains a nuclear localization signal (KKLKK^48^), it is likely that ATF4P3 retains its nuclear localization and DNA binding ability. Overall, *ATF4P3* is nearly completely intact (∼95%) at both the nucleotide (Fig 1A) and protein level (Fig. 1B-C).

#### ATF4P4

*ATF4P4* is located on Chromosome 11. Its uORFs 1 and 2 are extended due to changes in their stop codons (Fig. 1A). uORF1 now overlaps both uORF2 and uORF3. This uORF1 overlap likely results in inhibiting normal ribosomal binding to uORF3, which usually causes the CDS to be skipped^24–26^, and this likely would release some uORF inhibition of the *ATF4P4* CDS. ATF4P4 contains a premature stop codon that occurs after the p300 interaction domain, and this results in a 138 amino acid product. Despite this truncation, *ATF4P4* is nearly completely intact at the nucleotide level (∼95% identity). While ATF4P4 lacks both degradation domains and its DNA binding domain at the protein level, its p300 interaction domain remains intact (Fig. 1B-C). This suggests that ATF4P4 would lack targeted degradation while still maintaining its ability to bind p300. Within the 138 amino acid product (39.2% query coverage), ATF4P4 has 90.6% amino acid identity with ATF4 (Fig. 1).

Several sites of post-translational modification in ATF4 are changed in these retrocopies. ATF4P1/2 Long X has variants in two phosphorylation sites (T114A, T119S). These are two of four phosphorylated threonine residues, along with T107 and T115, important for protein stability^49^. Mutation of all four of these threonine residues causes a strong increase in ATF4 stability. It is possible that the T119S variant is still phosphorylated (as it is a serine). However, the T114A variant should remove the phosphorylation site, thereby increasing the stability of ATF4P1/2 Long X protein. ATF4P3 has a variant in a proline residue in the ODD domain (P162L), one of five prolines important for proper ODD domain-induced degradation^19^. This may lead to impaired degradation of ATF4P3 under normoxia, as this is observed for the same mutation in ATF4^19^. ATF4P4 has a variant in a phosphorylation site (S58F), and mutations in this same site in ATF4 lead to reduced transcription factor activity^50^. Given that ATF4P4 is missing its DNA binding domain, it is unclear what effect this variant amino acid has on ATF4P4. Overall, *ATF4P1-4* are relatively intact and have high nucleotide and protein identity with *ATF4*.

### Conserved and independent retrocopies of *ATF4* are common in mammals

If *ATF4* retrocopies are well-conserved across many species, it suggests they are under positive selection. As such, we examined the evolutionary history of each *ATF4* retrocopy. We analyzed *ATF4* retrocopies through manual BLAST-like Alignment Tool (BLAT) queries of 97 species, from yeast to humans, derived from UCSC-curated genome assemblies (Supp Table 2, University of California Santa Cruz (UCSC) genome browser^51^).

#### ATF4P2 and ATF4P1

Retrotransposition of *ATF4P2* occurred in a common ancestor of apes and *Cercopithecidae,* the monkeys of Africa and Asia, approximately 37.5 MYA (Fig. 2A, Supp Fig 2, Supp Table 2). In the six *Cercopithecidae* species we examined, *ATF4P2* contains the same ancestral early stop for its Long X protein, which results in a putative 131 amino acid product (Fig. 2A). This is different from what is observed in apes where they range from 85 to 170 AA in length. The only intact protein domain in *ATF4P2* Long X is the p300 binding domain.

**Figure 2.**
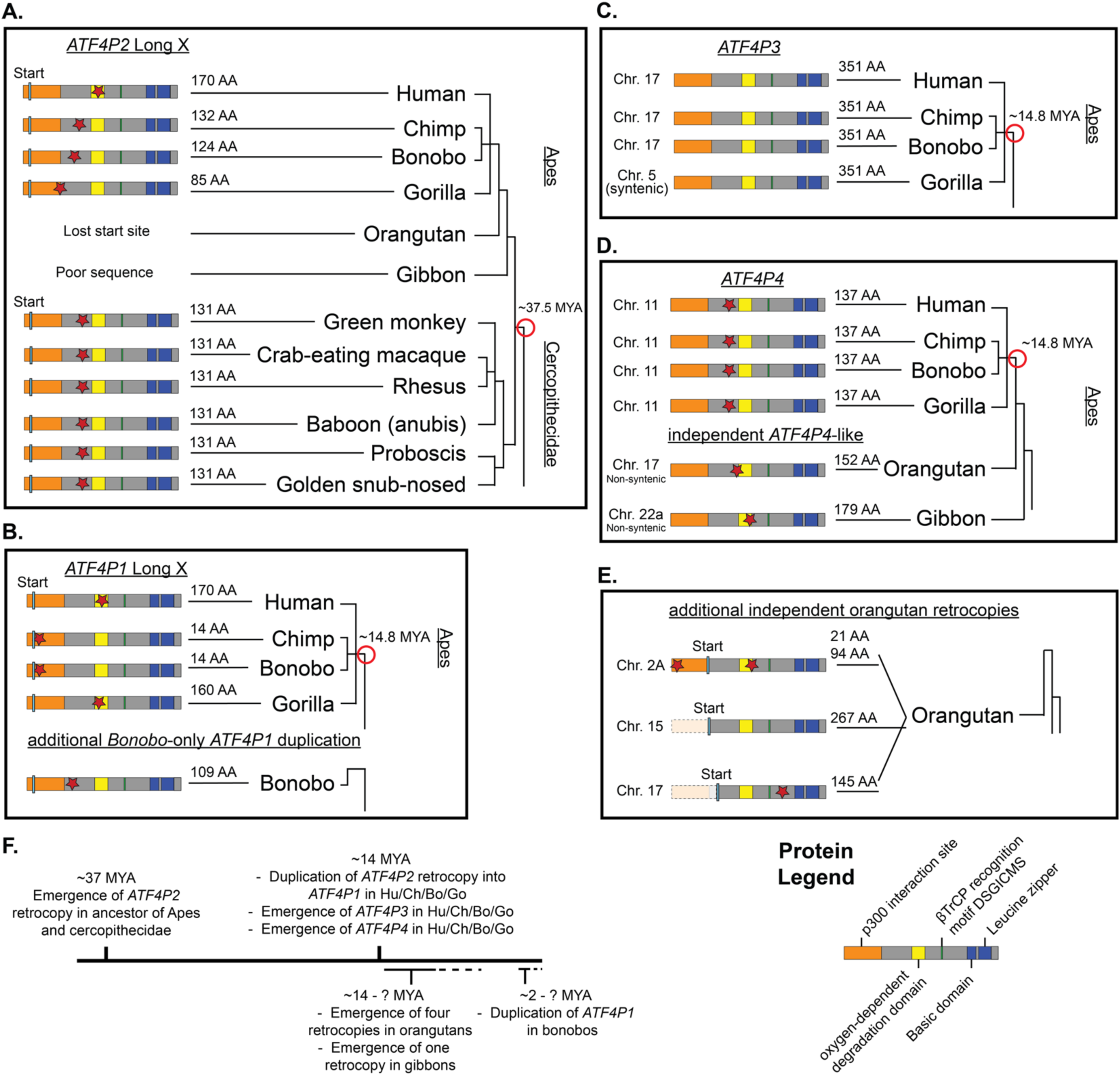
*Catarrhini* species tree comparisons of *ATF4* retrocopy and duplicate retrocopy proteins. **A.** *ATF4P2* arose ∼37.5 MYA in a common ancestor of *Catarrhini* (apes and *Cercopithecidae,* the monkeys of Africa and Asia). There is a short “Small X” peptide (ending at the blue bar) followed by a longer ORF (“Long X”) that resembles the truncated *ATF4P4*. Chimps and bonobos have lost “Long X”, but they maintain it in the duplication in *ATF4P1* and in an additional duplication in bonobos (see **B**). Note that orangutan only lost the start site of *ATF4P2* Long X, but they still maintain the highly truncated *ATF4P2* Small X (Supp Table 1). *Cercopithecidae* all contain the same truncating stop site resulting in a 131 amino acid product. **B.** *ATF4P1* resulted from a duplication of *ATF4P2* ∼14.8 MYA. In humans, it is an exact copy even at the nucleotide level, while chimps, bonobos, and gorillas have various levels of truncation. Bonobos also have an additional duplication of *ATF4P2* with its own unique truncation. **C.** *ATF4P3* arose in a common ancestor of apes ∼14.8 MYA, and it is fully intact at the amino acid level in each species we examined. **D.** *ATF4P4* arose in a common ancestor of apes ∼14.8 MYA, and it carries an early stop that truncates the protein to 137 amino acids. There are two independent *ATF4* retrocopies in orangutans and gibbons with similar truncations to *ATF4P4* but are non-syntenic. **E.** Orangutans have three independent (non-syntenic) retrocopies of *ATF4*. Two are genomically truncated (Chrs. 15 and 17), and Chr. 2A carries two early stop codons. This Chr.17 retrocopy is likely a partial duplication of the “ATF4P4-like” orangutan retrocopy in (D) given its proximity. **F.** Timeline of approximate emergence of new retrocopies and their duplications. AA = amino acid, MYA = million years ago.

**Table 1.** GTEx expression metrics in each tissue type. See Supp Table 4 for details.

|  | *ATF4* | *ATF4P1* | *ATF4P2* | *ATF4P3* | *ATF4P4* |
| --- | --- | --- | --- | --- | --- |
| # $\geq 0.1$ TPM | 52 | 0 | 0 | 52 | 49 |
| # $\geq 0.5$ TPM | 52 | 0 | 0 | 13 | 5 |
| Min TPM | 98.38 | 0 | 0 | 0.12 | 0.084 |
| Max TPM | 806.1 | 0.047 | 0.049 | 0.88 | 2.3 |

*ATF4P1* is an ape-specific duplication of the *ATF4P2* retrocopy. It likely arose ∼14.8 MYA because it is present in humans, chimps, bonobos, and gorillas but not orangutans or gibbons^52–56^(Fig. 2B). Human and gorilla *ATF4P1* genes have a putative Long X protein of 170 and 160 AA, respectively. In chimps and bonobos, their *ATF4P1* Long X sequence is truncated with an early stop resulting in a putative 14 amino acid long product (different from Small X protein). Bonobos also have an additional retrocopy duplication in tandem with its *ATF4P2* and *ATF4P1* (Fig 2B, Supp Fig 2, Supp Tables 1 and 3), resulting in three total copies. This additional bonobo duplication has 98% nucleotide identity with *ATF4P2*, and only 91% with *ATF4P1*, so it most likely also arose from *ATF4P2* sometime in the last ∼2 MY.

Human, chimps, and bonobos have a 30-34 bp internal duplication in *ATF4P2* and *ATF4P1* between the p300 interaction and ODD domains (Supp Table 1). In humans, this is exactly 30 bp (and thereby in-frame) in *ATF4P2* and *ATF4P1*, causing its Long X protein to be ten amino acids longer. However, this segment is 32 bp long in chimp *ATF4P2* and results in an early stop. The duplication likely has no impact on bonobo *ATF4P2* or *ATF4P1* or its additional duplication, or chimp *ATF4P1*, as other indels cause earlier stop codons. Unexpectedly, this ∼30bp duplication is not present in gorilla *ATF4P2* nor *ATF4P1* (2019 and 2016 genome assemblies via UCSC). As it is not present in orangutans or *Cercopithecidae* either, it is unlikely due to a deletion event in gorillas, and it is instead likely a new duplication arising after the gorilla split. However, because it is present in *ATF4P2* and *ATF4P1* (and the additional duplication in bonobos), this suggests the unlikely occurrence of two independent duplication events of this segment in *ATF4P2* and *ATF4P1* in the common ancestor of humans, chimps, and bonobos.

Given the divergence time of 14.8 MY, and a human mutation rate of ∼1.1 x 10^−8^/generation^57^, we expected to find ∼9 nucleotide changes between *ATF4P2* and *ATF4P1*. However, human *ATF4P2* is identical to *ATF4P1* at the nucleotide level. As described above, this region of the genome is highly invariant after a larger genomic duplication event^47^, suggesting that selection on something in this region is maintaining identity between the duplicates. Strikingly, this is not the case for the other apes examined, as human *ATF4P2* and *ATF4P1* are the only unchanged pair. Chimps, bonobos, and gorillas all have nucleotide differences between *ATF4P2* and *ATF4P1*, and the third copy in bonobos is also different from its *ATF4P2* and *ATF4P1* (Fig. 2A-B, Supp Table 1).

Of note, chimps and bonobos are very closely related species^58^, yet their *ATF4P2* and *ATF4P1* retrocopies contain several nucleotide and protein differences relative to each other. For example, while both have a putative truncated 14 amino acid long product in *ATF4P1*, these were independent, convergent events (22Gdel in bonobos and 5Cdel in chimps). In addition, ATF4P2 is truncated at a different amino acid length due to various nucleotide differences between chimps and bonobos (Fig 2A, Supp Table 1).

#### ATF4P3

*ATF4P3* arose in a common ancestor of humans, chimps, bonobos, and gorillas (Fig. 2C, Supp Fig 2, Supp Table 2). In each species, *ATF4P3* has a full protein length ORF, though each species has differences in amino acid variants (Supp Fig 3). Similar to humans, there are no changes to the DNA binding domain of ATF4P3 in chimps compared to ATF4. Bonobo ATF4P3 has one variant (K315Q) in its DNA binding domain, while gorilla ATF4P3 has two variants (R296C, N317S). We queried how these amino acid variants might impact protein function by using PolyPhen^59^ and SIFT^60^. K315Q in bonobo is predicted to be “benign” and “tolerated” by PolyPhen and SIFT, respectively. For the gorilla variants, both are predicted as “probably damaging” and “affecting protein function” by PolyPhen and SIFT, respectively. Given its sequence is identical in humans and chimps, and that the bonobo variant is predicted as benign/tolerated, this domain is likely functional if it is translated.

#### ATF4P4

Similar to *ATF4P3*, *ATF4P4* arose in a common ancestor of humans, chimps, bonobos, and gorillas (Fig. 2D, Supp Fig 2, Supp Table 2). In all species, ATF4P4 retrocopies are truncated by the same early stop codon, resulting in a 137 amino acid product. We also found two similarly truncated (but independent/non-syntenic) “*ATF4P4*-like” retrocopies in both orangutans and gibbons, creating 152 and 179 amino acid products, respectively (Fig. 2D). Thus, this truncated variant of an ATF4 retrocopy has arisen at least three independent times. Similar to ATF4P1/2, these truncated retrocopies only contain an intact p300 binding domain. Taken together, humans, bonobos, and gorillas all have three intact ATF4 retrocopies containing only the p300 binding domain (chimps have two). If expressed, this suggests some benefit to having extra copies of a p300-binding peptide.

#### Independent orangutan ATF4 retrocopies

Orangutans also contain three additional *ATF4* retrocopies that are non-syntenic to any other retrocopy we observed (Fig. 2E, Supp Fig 2, Supp Table 2). These retrocopies are unlike *ATF4P1-P4*. The Chr. 2A retrocopy has multiple internal start and stop sites, resulting in potentially smaller peptides. This results in one putative peptide that resembles *ATF4P1/2* Small X, but none that contain as much of the p300 domain as *ATF4P1/2* Long X. The retrocopy on Chr. 15 lacks the p300 domain due to a chromosomal truncation of DNA (unlike the previous protein-level truncations). The retrocopy on Chr. 17 may be a duplication of the “ATF4P4-like” retrocopy because it is only ∼1 Mb away. In addition, it is only a truncated copy of this gene with only one flanking tandem duplication site (TDS), rather than the two TDS sites expected if it were another LINE-1 element-derived retrocopy.

#### Approximate evolutionary timeline

The first retrotransposition event occurred ∼37 MYA (Fig. 2F). Later, approximately 14.8 million years ago (MYA)^52–56^, the common ancestor of humans, chimps, bonobos, and gorillas had three independent duplications of *ATF4* - two retroduplications (*ATF4P3*, *ATF4P4*) and one duplication of a retrocopy (*ATF4P1* duplicated from *ATF4P2*)^47^. This suggests a period of stronger selection for *ATF4* retrocopies ∼14.8 MYA (Fig. 2F). In apes, additional *ATF4* retrocopies and duplications occurred since then in orangutans (∼14.8 MYA or earlier) and bonobos (∼2 MYA or earlier) (Fig. 2F).

### Independent *ATF4* retrocopies arose in many other mammals

There are many genomically intact (e.g. not heavily degraded nor fragmented) retrocopies of *ATF4* throughout mammals. In addition to the extra retrocopies in orangutan and gibbon genomes (Supp Table 1-2, Supp Fig 2), there are multiple non-syntenic *ATF4* retrocopies in several other primates, including the Philippine tarsier (*Carlito syrichta*, four retrocopies), the mouse lemur (*Microcebus murinus*, one retrocopy), the bushbaby (*Otolemur garnetti*, three retrocopies), and the flying lemur (*Galeopterus variegatus*, three retrocopies) (Supp Fig 2, Supp Table 2). As none of these retrocopies are syntenic, there are at least twelve additional independent retroduplications of *ATF4* in primates, bringing the total we observed to eighteen (including the three duplications of retrocopies, Supp Fig 2). We also found evidence of independent duplications and retrocopies of *ATF4* in thirteen other mammals (Supp Fig 2, Supp Table 2). Most of these species only had a single retrocopy of *ATF4*, with the exception of the European hedgehog (*Erinaceus europaeus*, three retrocopies) and the nine-banded armadillo (*Dasypus novemcinctus*, four retrocopies). Of note, the *Bovidae* family has a syntenic retroduplication of *ATF4* present in at least bison, cows, and sheep (*Bison bison*, *Bos taurus*, and *Ovis aries*) (Supp Fig 2, Supp Table 2). This retrocopy is inserted within an intron of the *BANK1* gene, so it may utilize its nearby regulatory sequences. We see no evidence of *ATF4* retrocopies in birds, lizards, amphibians, fish, or any invertebrate species (using invertebrate orthologs, Supp Table 2). This is unsurprising, as there are much lower rates of retrocopies in non-mammalian species^61,62^.

### Likelihood of retaining ATF4 retrocopies over time

There are an estimated ∼8,000 retrocopies in humans^61,63^. Because many retrocopies arise with no nearby regulatory components, with only <20% having an intact ORF, it is not uncommon for neutral evolution to eventually degrade retrocopies^61,63^. Supporting this, we found several *ATF4* retrocopies in vertebrates that appear to have degraded over time (Supp Table 2). In contrast, persistence of an intact gene over time can suggest positive selection of that gene. *ATF4P1-4* are relatively evolutionarily young (within ∼37.5 MY, for example instead of a 90+ MY divergence between humans and mice^64^). This is also amplified by the long generation times in humans (∼25 years) and monkeys (∼10 years). Thus, it is possible that not enough time has passed to determine the likelihood of neutral evolution. However, given the multiple intact or conserved *ATF4* retrocopies in multiple ape and monkey branches, we wanted to computationally determine the likelihood of *ATF4P1-P4* ORFs persisting over time to date.

We first created consensus sequences of each retrocopy for *ATF4P1-P4* before modeling mutations over time (Supp Table 3). We then modeled the likelihood of maintaining the size of each retrocopy’s longest ORF over 10,000 simulations of decay within each species. Both *ATF4P1/2* and *ATF4P4* have truncated ORFs, and this short size makes them statistically more likely to avoid degradation. For *ATF4P1/2*, there was an approximately 45% chance of its Long X ORF remaining intact among apes and monkeys (Supp Fig 4A). For *ATF4P4*, with its much shorter evolutionary time, there was approximately a 97.6% chance of remaining intact (Supp Fig 4B). *ATF4P3*, on the other hand, is completely intact after the same evolutionary time period. As such, our model found approximately only a 20% chance of remaining intact within apes (Supp Fig 4B). While evolutionarily young, the chances do not favor keeping an intact ORF the size of *ATF4P3* in four different apes. As ATF4 is a transcription factor, we also wanted to model the likelihood of the DNA binding domain remaining completely intact in *ATF4P3* (Fig. 1B, Supp Fig 3). We determined the likelihood that this region remained intact with no non-synonymous changes or indels in humans and chimps. The DNA binding domain was unchanged at the protein level approximately 12.7% of the time, indicating this is even less likely to occur (Supp Fig 4C). This is also an underestimation, as our modeling does not take into account times when an upstream frameshift or nonsense mutation would remove the DNA binding domain. Overall, our modeling suggests at least *ATF4P3* may be under positive selection.

Lastly, we modeled the likelihood of human *ATF4P2* and *ATF4P1* remaining identical over 14.8 MY. As mentioned, this area of the genome is highly invariant^47^, so it is possible they are genetically linked to a different gene under positive selection. Nevertheless, it is interesting to model because as mentioned, unlike humans, the three other apes we examined do have changes between their *ATF4P2* and *ATF4P1* retrocopy duplications (Fig. 2A-B [Chimps, Bonobos, Gorillas], Supp Table 1). We used the same simulation as above, but we instead determined how many times there were zero changes at the nucleotide level. After modeling just ∼10 MY, all 10,000 simulated replicates had at least one nucleotide change (Supp Fig. 4D). As this duplication likely occurred ∼14.8 MYA^52–56^, this means there is a <0.01% chance of the *ATF4P2* duplication having no changes.

### *ATF4* retrocopies are expressed in tissues and human cells

To determine if the *ATF4* retrocopies are expressed in humans, we first queried the Genotype-Tissue Expression (GTEx) online portal for mRNA expression data of each retrocopy based on thousands of healthy individuals. *ATF4* has median transcript per million (TPM) values ranging from 98-806 depending on tissue (Supp Table 4, Table 1). In contrast, *ATF4P2* and *ATF4P1* have a maximum TPM value of 0.05, and the highest *ATF4P3* and *ATF4P4* median TPM values are 0.88 and 2.3, respectively (Table 1). GTEx uses a TPM cut-off of 0.1 to be considered expressed for use in their eQTL Analysis^65^, and the EMBL-EBI Expression Atlas sets a TPM cut-off of 0.5 for a gene to be considered expressed^66,67^. *ATF4P3* has a TPM of ≥0.1 in all tissues, with 13 at ≥0.5 TPM. *ATF4P4* has a TPM of ≥0.1 in 49/52 tissues, with 5 at ≥0.5 TPM. However, *ATF4P2* and *ATF4P1* never reach these thresholds. Overall, this suggests a low level of documented basal expression for *ATF4P3* and *ATF4P4* with none for *ATF4P2* and *ATF4P1*.

We next examined transcriptional markers around each retrocopy by using psiCube, which analyzes pseudogenes across humans, flies, and worms^68^. Based on psiCube annotation, both *ATF4P3* and *ATF4P4* have active chromatin (Pseudogene activity resource, psiCube, April 2014^68^). We examined this further by querying ENCODE data from seven human cell lines for chromatin markers^69^. *ATF4P3* and *ATF4P4* each harbor H3K4Me3 markers which are often found near transcription start sites^70^ (Supp Fig 5). We also note that *ATF4P4* is inserted directly next to a different retrocopy, *SH3GL1*, but the *SH3GL1* retrocopy has no H3K4Me3 markers (Supp Fig 5) - indicating specificity to *ATF4P4*. These active chromatin markers agree with the GTEx expression data, further lending support that these retrocopies are expressed in humans.

To verify the mRNA expression of these retrocopies, we tested endogenous retrocopy expression levels in a human cell line. Because *ATF4* interacts with tumor cell biology^7,11–16^, we avoided using common cancer-derived cell lines such as HeLa cells. Thus, we used hTERT RPE-1 cells, as they are a noncancerous human cell line that is karyotypically normal (CRL-4000®, ATCC)^71,72^. The *ATF4* retrocopies are highly identical to *ATF4* (Fig. 1A), so we designed primers to minimize off-target binding to each other. As the 3’ end of each primer is most important for binding and subsequent synthesis by polymerase^73^, we designed primers with a mismatch at the 3’ end of either the forward or reverse primer (see Methods). For *ATF4P1/P2*, we instead designed a primer around a large deletion only present in *ATF4P1/2*. Each ∼100-200 bp product contained single nucleotide variants (SNVs) with which we verified amplification of the correct cDNA. Importantly, we used DNAse treatment to eliminate genomic DNA contamination from our RNA samples, and we verified the efficacy of this treatment (Supp Fig 6). Using these primers and methodology, we sequenced RNA-derived cDNA from wild type RPE-1 cells.

Using Sanger sequencing and chromatogram analysis, we successfully identified expression of each specific retrocopy of *ATF4* (Fig. 3A). *ATF4P1/P2* and *ATF4P4* produced clean SNVs distinguishing them from both *ATF4* and the other *ATF4* retrocopies (Fig. 3A). However, *ATF4P3* was partially contaminated by *ATF4*, likely due to its high similarity to *ATF4* (>95%) preventing improvements to primer design. Nevertheless, we identified distinct *ATF4P3* transcripts via *ATF4P3*-specific nucleotide changes (Fig. 3A). Of note, we find that *ATF4P1/2* is indeed transcribed, despite its low expression on GTEx (Table 1). Surprisingly, in our subsequent qPCR analysis comparisons, *ATF4P1/2* has significantly more expression (at least double) than *ATF4P3* or *ATF4P4* (Supp Table 5). This difference may be due to cell type specificity, or that our PCR method would pool copies of *ATF4P2* and *ATF4P1* together. Regardless, *ATF4P1/2* can have much higher mRNA expression than databases suggest, which increases the likelihood of its biological significance. Overall, we find that all *ATF4* retrocopies are endogenously transcribed in RPE-1 human cells.

**Figure 3.**
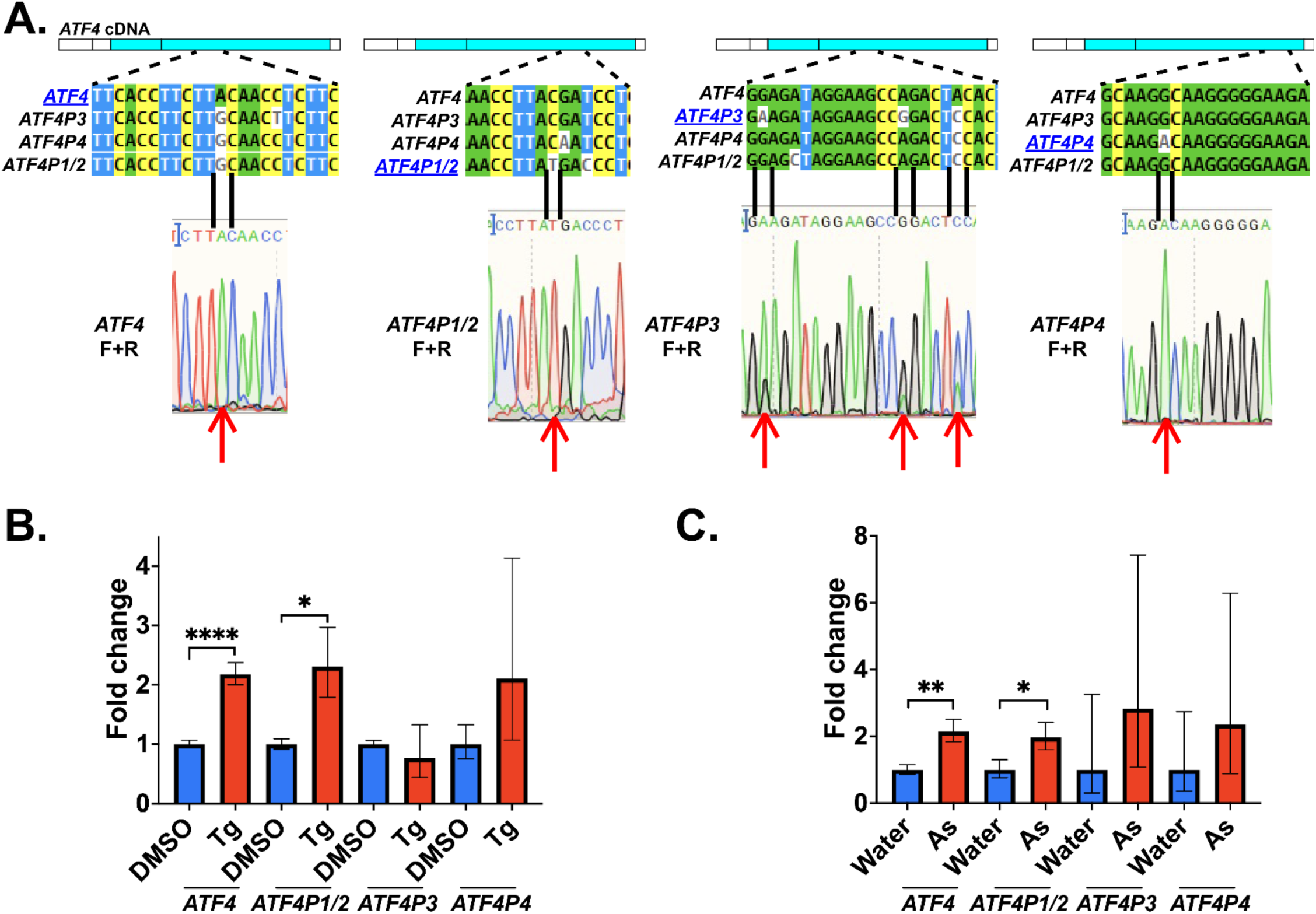
Chromatogram analysis identifies endogenous *ATF4* retrocopy transcripts in human cells. **A.** We used primers specific to each gene to amplify transcripts under normal conditions. Using SNPs, we identified sections of each amplified transcript that differed between each gene. Black bars denote the SNPs used, and the transcript being examined is in blue. We checked the chromatograms of these regions to determine how “clean” each transcript signal was (e.g. whether *ATF4*, or other retrocopies, were being partially amplified). *ATF4*, *ATF4P1*/*2*, and *ATF4P4* had clean peaks, while *ATF4P3* had slight contamination from *ATF4* transcripts. Overall, we were able to detect that every retrocopy transcript was present in cells. We created alignments using ClustalOmega and visualized them using MView^74^. **B.** Using the primers from (A), thapsigargin (Tg) treatment induced *ATF4* and *ATF4P1/2* expression (N=5 biological replicates, ****p<0.0001, *p<0.05, Student’s t test). **C.** Like Tg treatment, treatment with arsenite (As) induced *ATF4* and *ATF4P1/2* expression (N=3 biological replicates, **p<0.01, *p<0.05, Student’s t test). For **B-C**, we plot the average fold change and the standard error.

*ATF4* expression is transcriptionally and translationally induced by multiple inputs into the integrated stress response (ISR)^1^. We tested whether inducing the ISR also induced mRNA expression of *ATF4* retrocopies. We exposed wild type RPE-1 cells to thapsigargin (TG), a SERCA inhibitor that causes ER stress and activates the PERK branch of the ISR^1,75^. As previously shown^76–78^, TG induces mRNA expression of *ATF4* (Fig. 3B). Remarkably, TG also increased mRNA expression of the retrocopy *ATF4P1/2*, to approximately the same fold increase (∼2-fold). TG did not significantly increase *ATF4P3* or *ATF4P4*. To determine if the *ATF4P1/2* increase in expression was from ISR induction, and not a TG-specific response, we tested a second inducer of the ISR. We exposed cells to sodium arsenite - an inorganic compound that activates the HRI branch of the ISR^1,79^. As with TG, arsenite induced mRNA expression of both *ATF4* and *ATF4P1/2* (Fig. 3C), but it did not significantly increase *ATF4P3* or *ATF4P4*.

While neither ISR-inducing drug significantly increased *ATF4P3* nor *ATF4P4* expression, we observed that some replicates showed strong responses to treatment, while other replicates showed a smaller effect (Supp Table 5). Here, *ATF4* expression appeared to be correlated to retrogene expression. For example, in one TG-treated cell sample, *ATF4* had at most a 2.6x increase in expression, and the same cell sample increased *ATF4P1/2* by 5.5x, *ATF4P3* by 5x, and *ATF4P4* by 16.2x (Supp Table 5). Under arsenite treatment, the cell population replicate with the strongest increase in *ATF4* expression was 2.7x, while the same cell population increased *ATF4P1/2* by 2.3x, *ATF4P3* by 6.9x, and *ATF4P4* by 8.2x (Supp Table 5). This suggests that *ATF4P3* and *ATF4P4* can in fact be induced by the ISR, but they may require more precise timing or dosage to fully capture how their expression is affected. Overall, *ATF4P1/2* mRNA expression is induced by ISR signaling to approximately the same fold change as *ATF4*, while *ATF4P3* and *ATF4P4* mRNA expression can be induced only under certain ISR timings or conditions.

### *ATF4* retrocopies can be translated and are degraded by the proteasome

To determine the effects of these retrocopies in human cells, we used lentiviral transfection to create stable overexpression (OE) lines of *ATF4* and each of the *ATF4* retrocopy ORFs in RPE-1 cells. We created each OE cell line and confirmed correct sequencing - including the presence of an in-frame V5 tag. Using the V5 tag, we verified correct band sizes for each of the overexpressed proteins except for the ATF4P1/2 Small X retrocopy, from which we were unable to verify protein on our Western blots (likely due to its small size) (Supp Fig 7A-B). These OE cell lines had no gross defects when compared to wild type cells.

As described, ATF4 has two degradation domains: 1. an oxygen-dependent degradation (ODD) domain; and 2. a ßTrCP motif. Each of these domains result in the sequestration to, and termination at, the proteasome^1,19,23^. As such, inhibiting the proteasome stabilizes ATF4 protein levels^23,80^, so we tested if that is also true for ATF4 retrocopies. To inhibit the proteasome, we used bortezomib (Btz)^80,81^. As expected, ATF4 protein increased under Btz treatment (Fig. 4A, Supp Fig 7B-C). In each retrocopy OE cell line, their protein was also increased under proteasome inhibition (Fig. 4A, Supp Fig 7B-C), indicating that these are also degraded by the proteasome.

**Figure 4.**
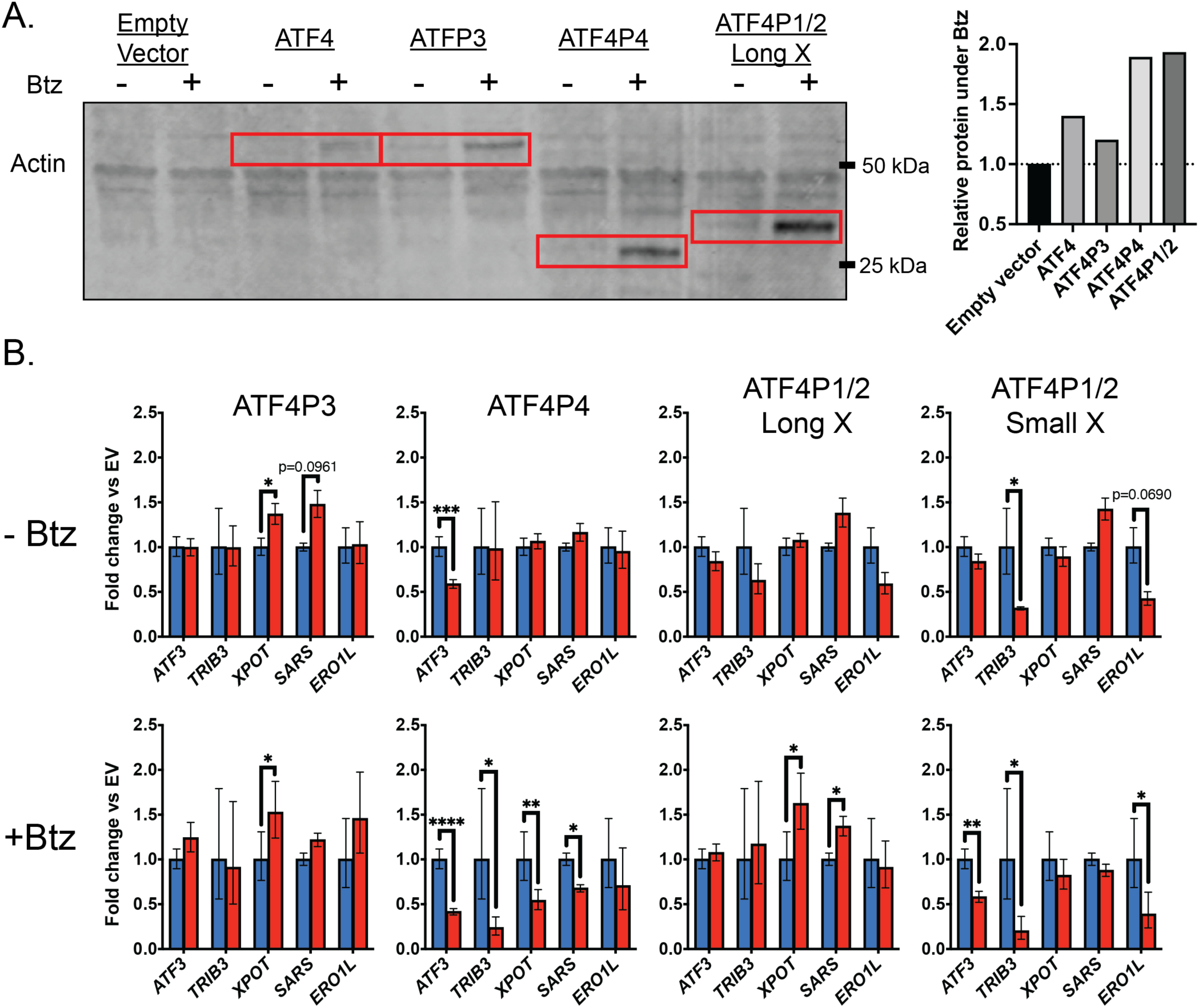
*ATF4* retrocopy proteins are degraded by the proteasome and affect the expression of canonical *ATF4* gene targets. **A.** We grew each cell line overexpressing *ATF4* and its retrocopies with or without proteasome inhibition (through Btz). Proteasome inhibition increased the ratio of *ATF4*/retrocopy protein in each instance (see red boxes and rightmost graph). Note that image contrast was globally adjusted for band clarity - see full gel in Supp Fig 7. **B.** We analyzed gene expression of five known *ATF4* target genes in each overexpression cell line, with or without proteasome inhibition. Generally, the increased levels of each retrocopy protein from proteasome inhibition (in **A**) had a stronger impact on the expression of each *ATF4* target gene. However, there were gene expression changes under both conditions, indicating that these retrocopies can impact *ATF4* gene expression (N=3 biological replicates, ****p<0.0001, ***p<0.001, **p<0.01, *p<0.05, One-way ANOVA with adjusted p-values derived using Dunnett’s multiple comparison correction).

Surprisingly, while ATF4P4 and ATF4P1/2 lack both of the ATF4 degradation domains (Fig. 1), they still showed increased protein levels with proteasome inhibition. We also observed a second band at ∼10-12kD in the ATF4P4 OE line (Supp Fig 7A-C) that also increased under Btz treatment. We hypothesize that this product is from a second methionine start site within ATF4P4 (annotated in Supp Fig 7D). Because this smaller product of ATF4P4 also increased under Btz treatment, it is likely also degraded by the proteasome. As such, we looked within this product for potential lysine residues with evidence of ubiquitination in ATF4 (using Uniprot^82^ - P18848). ATF4 residue K88 has high confidence as a ubiquinated site^82,83^, and it is maintained in ATF4P4 and ATF4P1/2 Long X (Supp Table 1). Thus, we hypothesize that ubiquitination of residue K88 might affect its proteasomal degradation, as it would explain why ATF4P4 and ATF4P1/2 have higher abundance under Btz treatment despite lacking the ODD domain and ßTrCP motif. Finally, we also note that ATF4P4 and ATF4P1/2 have a stronger relative response to Btz treatment (Fig. 4A, Supp Fig 7B-C), suggesting they have a faster turnover rate by the proteasome. The proteasome must unfold ubiquitinated proteins prior to processing^84^. Given that ATF4P4 and ATF4P1/2 are both truncated, they may have a less stable folded product and are thereby more easily processed by the proteasome. Taken together, all ATF4 retrocopy proteins are degraded by the proteasome, similar to ATF4.

### *ATF4* retrocopies affect expression of canonical *ATF4* target genes

ATF4 is a transcription factor that affects the expression of hundreds of genes^4–6^. We wanted to determine what effect, if any, ATF4P1-4 has on the expression of a diverse set of ATF4 target genes. We selected five ATF4 transcriptional target genes relating to unique downstream pathways: *TRIB3*, involved in negative feedback of ATF4^1^; *ATF3*, a transcription factor important for stress and regeneration^1^; *SARS*, a tRNA charging protein^5^; *XPOT*, involved in export of tRNAs^5^; and *ERO1L*, involved in protein folding^5^. We determined the expression of these genes under ATF4 and retrocopy overexpression. To simulate cell stress resulting in increased expression of these proteins, we also determined gene expression under Btz treatment. Note that we included our nucleotide sequence-verified ATF4P1/2 Small X cell line in this assay, though we could not verify the expression of its small peptide (Supp Fig 7B).

ATF4P3 increased *XPOT* expression, with or without Btz treatment (Fig. 4B). It also increased *SARS* expression without Btz and *ERO1L* expression with Btz, but neither to a statistically significant level. As ATF4P3 most closely resembles ATF4, we expected that this protein would be most able to increase expression of ATF4 target genes. However, to our surprise, overexpression of ATF4 did not increase expression of these target genes, which suggests that either ATF4 levels are already high in our cells, or that tight regulation of ATF4 protein degradation or translocation prevents any further effect on gene expression. If it is the latter case, it is especially interesting considering that ATF4P3 is capable of increasing gene expression - suggesting one or more of its fifteen single amino acid variants (Fig. 1) can evade this tight regulation.

ATF4P4 had the strongest overall effect on the expression of ATF4 target genes, but surprisingly it decreased expression of those genes rather than increased them (Fig. 4B). ATF4P4 significantly reduced the expression of *ATF3* with or without Btz. ATF4P4 also reduced the expression of *TRIB3*, *XPOT*, and *SARS,* but only with Btz treatment. ATF4P4 lacks the basic leucine zipper domain, including its internal nuclear localization signal (Fig. 1), so this is unlikely due to any DNA binding activity. The truncated form of ATF4P4 leaves only the p300 binding domain intact (Fig. 1B). Thus, one potential mechanism is that ATF4P4 competes with ATF4 for p300 binding, thereby reducing the stability of ATF4 and its transcription of DNA^18^. Alternatively, ATF4P4 may bind ATF4 co-binding proteins, such as CHOP^17^, to prevent formation of heterodimers necessary for gene transcription. Overall, ATF4P4 expression negatively impacts the expression of several canonical ATF4 target genes.

While ATF4P1/2 Long X has a similarly truncated protein like ATF4P4 (Fig. 1), it had a smaller effect on gene expression (Fig. 4B). Under Btz treatment, ATF4P1/2 Long X increased both *XPOT* and *SARS* expression. However, ATF4P1/2 Long X lacks the first 16 amino acids of the p300 binding domain, which may weaken any potential binding to p300 and any subsequent effects on ATF4. This is further supported by the fact that the ATF4P1/2 Small X peptide, which is simply the first 15 amino acids of the p300 binding domain (with one change, D15A) (Fig. 1B-C), mimics ATF4P4 by also having a negative effect on ATF4 target gene expression. With or without Btz treatment, ATF4P1/2 Small X reduced *TRIB3* expression (Fig. 4B). Under Btz treatment, ATF4P1/2 Small X reduced the expression of *ATF3* and *ERO1L*. ATF4P1/2 Small X also reduced expression of *ERO1L* without Btz treatment, but not to a statistically significant degree. Given these results and the amino acid overlap of ATF4P4 and ATF4P1/2 Small X, it is possible that the first 15 amino acids of the p300 interaction domain has a negative effect on ATF4 target gene expression.

Overall, it is clear that with or without cell stress, ATF4 retrocopies can alter ATF4 target gene expression. Given this, their similar protein degradation patterns (Fig. 4), their ability to be upregulated by the ISR (Fig. 3), and their expression in humans (Fig. 3), we conclude that *ATF4P1-4* are functional retrogenes.

## Discussion

We characterized four retrogenes of the stress transcription factor, *ATF4*, in humans. We found these retrogenes across multiple primate lineages, suggesting an increase in their selection ∼14.8 MYA. These retrogenes are expressed at the mRNA level in cells - including *ATF4P1/2,* which responds to induction of the integrated stress response (ISR). Overexpressing stable proteins of *ATF4* retrocopies alters the expression of *ATF4* target genes, indicating their potential for biological function. Given the important role for ATF4 in human diseases, such as cancer^7,11–16^, our findings suggest that these *ATF4* retrogenes should be considered when studying the ISR and ATF4 pathways.

The increase in LINE-1 element transposons in mammalian lineages resulted in an increase in their total retrocopied genes^61,85^. Humans have ∼8,000 retrocopies, but most remain uncharacterized or unstudied^33,61,63,86^. However, there are several instances of retrocopies having biological function. The retrogene *retroCHMP3* inhibits viral budding^34,87^. The testis-specific retrogene *PGK2* positively regulates sperm function^39,88^. At the mRNA level, the retrogene *PTENP1* regulates PTEN expression at the mRNA level^35,36^. Its transcript acts as a decoy that pulls inhibiting microRNAs away from PTEN, thereby increasing its expression^35,36^. Thus, *ATF4P1-4* are yet another example of retrogenes capable of affecting cell biology.

Approximately 30% of retrocopies have RNA expression data^85,89^. However, previous expression data must be validated. For example, there is evidence of *ATF4P1/2* mRNA expression in our cells, yet *ATF4P1/2* does not have significant expression in GTEx^90^. Therefore, individual retrocopy expression should be more closely examined on a case-by-case basis. In particular, mining new long-read RNA-seq data for retrocopy sequences would likely be fruitful given the high percentage identity that retrocopies often have to their parental genes.

The ISR is a critical stress response pathway conserved from yeast to humans^1,91^. It is crucial for restoring cell homeostasis under various stressors^1^. It is dysregulated in some patients with Wolcott-Rallison syndrome^92^ and pulmonary disease^93^, as well as in several other disease models^94–96^. We found that the retrogenes *ATF4P1*/*2* are upregulated under the ISR stressors arsenite and thapsigargin^79,97^ (Fig. 3). In addition, overexpression of *ATF4P1/2* altered expression of several canonical *ATF4* target genes (Fig. 4) - positively or negatively depending on the ORF involved. The *ATF4P1/2* retrogene is the oldest retrocopy in humans (Fig. 2) and both copies remain unchanged at the nucleotide level for ∼14 MY in humans. Thus, *ATF4P1/2* may actively participate in the ISR in humans and should be considered when further studying these pathways and disease models. *ATF4P3* and *ATF4P4* could also be induced under ISR stress (Fig. 3), but they did have more replicate variability which may indicate different conditions for their expression. For example, *ATF4P3* is inserted in the intron of *RNF157* (Supp Fig 1) which has strong brain expression (^98^ and also via GTEx^90^). Thus, it may require a neuronal model or cell line to fully capture the expression or effects of *ATF4P3*.

Retrocopies can affect the expression of their parent genes at the transcriptional or translational level^33–36,89^. Based on our proteasome inhibition experiments (Fig. 4), all four *ATF4* retrogene proteins are typically degraded by the proteasome, like ATF4. Inhibiting the proteasome, and thus increasing their protein levels, alters the expression of ATF4 target genes. This suggests that their actual translated peptides can influence gene expression. While this could be explained by an intact DNA-binding domain in ATF4P3, it does not explain ATF4P4 or ATF4P1/2 which lack this domain. ATF4 typically forms a heterodimer, such as with CHOP or ATF3^17^, to bind to DNA. It is possible that ATF4P4 or ATF4P1/2 can bind - even partially - to these co-binding partners and sequester them away from ATF4. This could reduce gene expression overall or force *ATF4* to bind with different co-binding partners and switch which genes are upregulated. Alternatively, it is possible that these retrogenes act at the RNA level (like retrogene PTENP1^35,36^) - as our overexpression clones would presumably have increased mRNA as well - or a combination of the two. The exact mechanism of how these retrogenes influence *ATF4*, and its transcriptional ability, will be of interest to further characterize how they impact the ISR and related diseases.

While they lack their DNA binding domain, ATF4P4 and ATF4P1/2 maintain their p300 interaction domain (Fig. 1). As p300 usually stabilizes ATF4 and helps prevent its degradation^18^, it is possible that ATF4P4 and ATF4P1/2 sequester p300 away from ATF4 and thereby reduce its expression. Additionally, the ability to bind p300 offers an explanation for the evolutionary origin of these retrogenes, including our finding that humans, chimps, bonobos, and gorillas have at least two “p300-only” retrocopies. Certain viruses can hijack p300 and use it to aid the transcription of viral machinery^99–101^. For example, the HIV-1 Tat protein is acetylated by p300 to aid its transcription^102^, and adenovirus E1A uses p300 to modulate gene expression^103^. Outside of any effect on ATF4 target gene expression, it is possible that ATF4P4 and ATF4P1/2 sequester p300 away from viruses to help limit viral gene transcription. Given that the ISR is known to have evolutionary pressure from viruses^40,41^, this provides a potential source of evolutionary pressure and selection to help explain the presence of maintained, but truncated, *ATF4* retrogenes.

## Materials and Methods

### Databases accessed for gene queries, expression, and protein folding

We used the UCSC genome browser to identify retrocopies of *ATF4* by using BLAT^51^ to search using the species-specific endogenous parental *ATF4* gene. We identified retrocopies, with a BLAT matching score of at least 100, by the presence of a PolyA tail and/or by a lack of introns. We noted cases of uncertainty, such as with short contigs or poor (e.g. “N”-rich) sequencing around potential retrocopies. In species where *ATF4* was not annotated, we used a closely related species (when possible) to predict the parental *ATF4* location in the queried species, then used that section to search that species for potential retrocopies. We also noted any duplications of *ATF4* in Supp Table 2, but only for annotation purposes, and we do not discuss them here.

We annotated exons and introns of human *ATF4* by referencing the Ensembl genome database^104^. We specifically used the canonical *ATF4* transcript ENST00000674920.3 (ATF4-206). We referenced the UniProt database to confirm PTM sites^82^. We used MacVector™ (v.18.2.5.) to help annotate and visualize DNA sequences.

We predicted protein structures using AlphaFold 3 using default settings^105^. We downloaded .CIF files of each protein and oriented them using PyMol (The PyMOL Molecular Graphics System, Version 2.2.6 Schrödinger, LLC.). To help visualize their structures, we used PyMol to color specific amino acids for Figure 1.

To create our species tree, we first generated our .tre file using the “rotl” package in Rstudio (version. 2023.12.1+402, RStudio Team (2020). RStudio: Integrated Development for R. RStudio, PBC, Boston, MA URL http://www.rstudio.com/). We uploaded this file to the interactive Tree of Life^106^ to correct minor labeling errors and to sort species.

### Cells

RPE-1 cells (hTERT RPE-1, CRL-4000, ATCC) were a gift from Drs. Michael Boland and David Goldstein (Columbia University). HEK293T cells were gift from Dr. Charles Murtaugh (University of Utah).

We maintained RPE-1 cells in DMEM/F-12 media (Gibco™, ThermoFisher cat. 11320033) supplemented with 10% FBS (Gibco™, ThermoFisher cat. 26140079) and 0.01 mg/ml hygromycin B (Sigma-Aldrich, cat. H3274, diluted in water). We maintained HEK293T and HeLa cells in DMEM with pyruvate (Gibco™, ThermoFisher cat. 11995065) supplemented with 10% FBS (Gibco™, ThermoFisher cat. 26140079). We grew both cells in cell incubators set to 37°C and 5% CO_2_.

### Determining ATF4 retrocopy expression via database and qPCR

We used the Genotype-Tissue Expression (GTEx) Project to determine expression levels of *ATF4* retrocopies (*ATF4P1-P4* are annotated on GTEx). GTEx was supported by the Common Fund of the Office of the Director of the National Institutes of Health, and by NCI, NHGRI, NHLBI, NIDA, NIMH, and NINDS. The data used for our analyses here were obtained from the GTEx Portal on under the dbGaP Accession phs000424.v10.p2 on 02/17/2025.

To design primers for *ATF4*/retrocopy expression, we first aligned *ATF4* and its retrocopies using ClustalOmega^107^. As we needed to use primers specific to each copy, without binding each other, we focused on DNA segments with low similarity between each retrocopy. When such areas were small, we used that low similarity area as the 3’ end of each primer, as this is the most important for primer binding/synthesis^73^ (see Primer Table). We used qPCR primers for *ATF4* target genes from PrimerBank^108^ or designed them using Primer3Plus^109^. Primers were ordered from IDT DNA (Coralville, Iowa).

To generate RNA, we first grew cells in 6-well plates until ∼80% confluent. We then added TRIzol™ Reagent (15596026, Invitrogen) to wells and scraped cell lysates into microcentrifuge tubes. We vortexed tubes for at least 15 seconds, and we stored them at −80°C for at least 24hrs before further processing. We used the DirectZol™ kit (Zymo Research, cat. R2052) to extract RNA, and the ProtoScript® II First Strand cDNA Synthesis Kit (NEB, cat. E6560L) to convert RNA to cDNA. We analyzed gene expression using ATF4/retrocopy-specific primers (Primer Table), PowerUp™ SYBR™ Green Master Mix (Fisher Scientific, cat. A25918), and QuantStudio 3 for analysis (ThermoFisher).

### ORF decay modeling

We aligned *ATF4* sequences using Multiple Sequence Alignment by Log-Expectation (MUSCLE 3.8)^74^ of either apes or *Catarrhini* to represent ancestral copies of 14.8 MYA and 37.5 MYA species, respectively. We then created estimates of ancestral *ATF4* by inputting these MUSCLE-aligned sequences into EMBOSS Cons^74^ to create consensus sequences using minimal manual annotation (see Supp Table 3).

We used the python script ‘mutator’ to simulate the decay of *ATF4* retrocopies under neutral evolution, and ‘orf_scanner’ to count open reading frames^110,111^. We modified the code of the ‘mutator’ script to use a Poisson distribution (rather than Gaussian) to better model the relatively short evolutionary times of these retrocopies (see new code at https://github.com/hansmdalton/pedestal_fork). As we are most focused on the human retrocopies, we used human substitution, insertion, and deletion rates as determined from the literature^111,112^: substitution = 1.16e-8, insertion = 2e-10, deletion = 5.5e-10. We used a generation time of 25 years for *ATF4P3* and *ATF4P4* to represent apes^113^, and we used a generation time of 10 years for *ATF4P1/P2* to represent monkeys within *Catarrhini* that still maintain this retrocopy^114–116^. We derived the estimates of 14.8 MYA divergence of orangutans and 37.5 MYA divergence of *Platyrrhini* from the literature^52–56^. We simulated ORF decay in 10,000 replicates every 50,000 years for 50,000,000 years.

Starting from our estimated ancestral copy (Supp Table 3) at 14.8 MY or 37.5 MY, we determined probabilities of each lineage-specific branch terminus retaining the intact retrocopy ORF being tested (e.g. full 351 AA for ATF4P3, or >130 AA for ATF4P1/2 Long X). To prevent double-counting branches, we sectioned the phylogenetic branches into non-overlapping segments for testing species probabilities. To determine the overall probability, we then multiplied these probabilities together for each ORF. For amino acid changes in the DNA binding domain of *ATF4P3*, we used Biopython^117^ to translate each permutation to look for any non-synonymous changes or indels. For looking for any nucleotide changes in *ATF4P2*, we ran the same mutator script as above, but then we counted (using Python) the presence of any nucleotide change or indels. We then plotted all of these results using Rstudio^118^ and annotated percentages and labels in Adobe Illustrator.

### Target Site Duplication Identification

The presence of poly-A sequences and the lack of introns within the sequences of retrocopies indicates that they likely arose from retrotransposons^33,87^. This is typically driven by LINE-1 elements which leave a duplicated sequence artifact flanking each retrocopy^33,87^. We manually assessed the ∼100 bp downstream of the poly-A sites following the 3’ UTR (as determined from parental *ATF4*). We used this sequence to identify identical sequences upstream of the 5’ UTR. When we identified a likely TSD site, we checked this TSD against the rest of the species for this same sequence, refining the sequence as needed.

### ISR stress treatments

We grew wild type RPE-1 cells to ∼80% confluency for the experimental treatment day. We treated cells with either 0.1 uM thapsigargin (Tg) (Sigma cat. S7400) for 8hrs (vehicle = DMSO), or 10 uM sodium arsenite (As) (Sigma cat. T9033) for 6hrs (vehicle = water). We plated 3-5 biological replicates of WT RPE-1 cells in 6-well plates (As) or 10cm plates (Tg). After treatment, we collected cells in TriZol and performed qPCR analysis as above.

### Lentivirus production and transfection

As *ATF4* retrocopies have high similarity to each other at the nucleotide level, we employed two sequential PCR reactions to specifically amplify each retrocopy for cloning (Primer Table). Due to differences in introns, the cloned sequence for *ATF4* was derived from RNA (cDNA), while the retrocopies were derived from genomic DNA. We derived genomic DNA from HeLa cells and RNA from human fibroblasts^119^ (gifts of Dr. Nels Elde, University of Utah). We converted RNA to cDNA using the ProtoScript® II First Strand cDNA Synthesis Kit (NEB, cat. E6560).

First, we used primers ∼100-500 bp up- and downstream of each *ATF4* retrocopy to amplify that region. We then gel extracted (QIAquick Gel Extraction Kit, cat. 28704) this product to purify it from other genomic DNA. We this as the product for further Gateway cloning using primers specific to each gene. Because the SmallX insert was so short (55 bp), we instead ordered this entire sequence from IDT DNA (Coralville, Iowa) ready for further Gateway cloning. We sequenced all inserts and final products to ensure these were correct (Sanger sequencing via the University of Utah HSC DNA Sequencing Core).

For all copies of *ATF4* (including the parental gene *ATF4* itself), we observed no protein expression when we directly cloned either the CDS alone or the 5’ UTR plus CDS (performed in both RPE-1 and HeLa cells). To help resolve this, we designed primers to modify the native Kozak sequence^120^ upstream of the first codon, while maintaining the coding sequence integrity. Using these modified Kozak sequences, we observed protein expression of all clones.

**Table 2.** Native *ATF4*/retrocopy Kozak sequences are relatively weak and required modification to induce correct protein expression in our cells.

| Gene | Endogenous Kozak | Modified Kozak |
| --- | --- | --- |
| *ATF4* | CGCAACATGACC | GCCACCATGACC |
| *ATF4P3* | CGCAACATGACC | GCCACCATGACC |
| *ATF4P4* | CGCAACATGACC | GCCACCATGACC |
| *ATF4P1/2*, Small X | TGCACCATGACC | GCCACCATGACC |
| *ATF4P1/2*, Long X | GGCTTGATGTCC | GCCACCATGTCC |

We used the Gateway cloning system (Invitrogen™, ThermoFisher cat. K240020) to generate the entry vectors used for creating ATF4 overexpression cell lines. We then used LR clonase (Invitrogen™, ThermoFisher cat. 11791020) to transfer our clones into a lentiviral plasmid containing a C-terminal V5 tag (Addgene, pLenti6.2-ccdB-3xFLAG-V5, Plasmid #87071). We used a C-terminal tag to avoid future conflicts with 5’ UTR/Kozak/uORF analyses. We then generated lentivirus particles in HEK293T cells via FuGENE® 6 Transfection Reagent (Promega cat. E2693) using psPAX2 and pCAG-VSVG packaging plasmids (Addgene plasmid #12259, and Addgene #35616, respectively. Gifts of Dr. Charles Murtaugh, University of Utah).

We incubated RPE-1 cells with our generated lentivirus particles along with 8 μg/ml polybrene (gift from Dr. Charles Murtaugh, University of Utah). After 48hrs, we selected for transfected polyclonal cell populations using 10 μg/ml blasticidin (Gibco™, ThermoFisher cat. R21001). We then selected for monoclonal populations by picking cell colonies to individual plates where we continued blasticidin selection and ensured single colony expansion. We then performed western blot analyses (see below) to check for accurate and robust protein production of each cloned retrocopy.

### Proteasome inhibition

To induce proteasome inhibition, we first grew RPE-1 cells to 80-90% confluency. We then replaced their regular media with media containing either 0nM or 100nM bortezomib (both with 0.1% DMSO). After six hours of exposure to bortezomib, we collected cell protein for Western blot or cells in TriZol for RNA extraction and downstream qPCR analysis.

### Western blots

To collect protein, we washed cells with PBS, then treated them with RIPA buffer supplemented with 1x Protease Inhibitor Cocktail (Cell Signaling Technologies, cat. #5871). We lysed cells for 30 minutes at 4°C under agitation, then centrifuged at 14,000g for 20 minutes. We extracted supernatants and determined protein concentration using the bicinchoninic acid (BCA) assay (Pierce™ BCA Protein Assay Kit, cat. 23227). We then froze samples at −80°C for later use.

To denature proteins, we added 6x loading buffer supplemented with β-Mercaptoethanol (Sigma, cat. M6250) and heated them at 95°C for 5 minutes. We resolved proteins on using a 12% Criterion™ XT Bis-Tris Protein Gel (Bio-Rad #3450118) and XT MES Running Buffer (Bio-Rad, cat. #1610789). We detected proteins using anti-V5 (Cell Signaling Technologies, cat. 13202, 1:1,000), anti-ATF4 (Proteintech, cat. 10835-1-AP, 1:800), anti-beta actin (Santa Cruz Biotechnology, cat. sc-47778, 1:200) primary antibodies, and anti-Rabbit and anti-Mouse secondaries (“DyLight”, Cell Signaling Technology cats. 5151 and 5470, respectively, 1:10,000). We used the Precision Plus Protein™ All Blue Standards for our ladder (Bio-Rad, cat. #1610373). We visualized blots using an Odyssey DLx imager (LI-COR).

### Statistics and graphs

We performed statistics and created figures using GraphPad Prism (version 10 for Windows, GraphPad Software, Boston, Massachusetts USA, www.graphpad.com). We also utilized Rstudio (version 2023.12.1+402, RStudio Team (2020). RStudio: Integrated Development for R. RStudio, PBC, Boston, MA URL http://www.rstudio.com/)

**Primer table.**
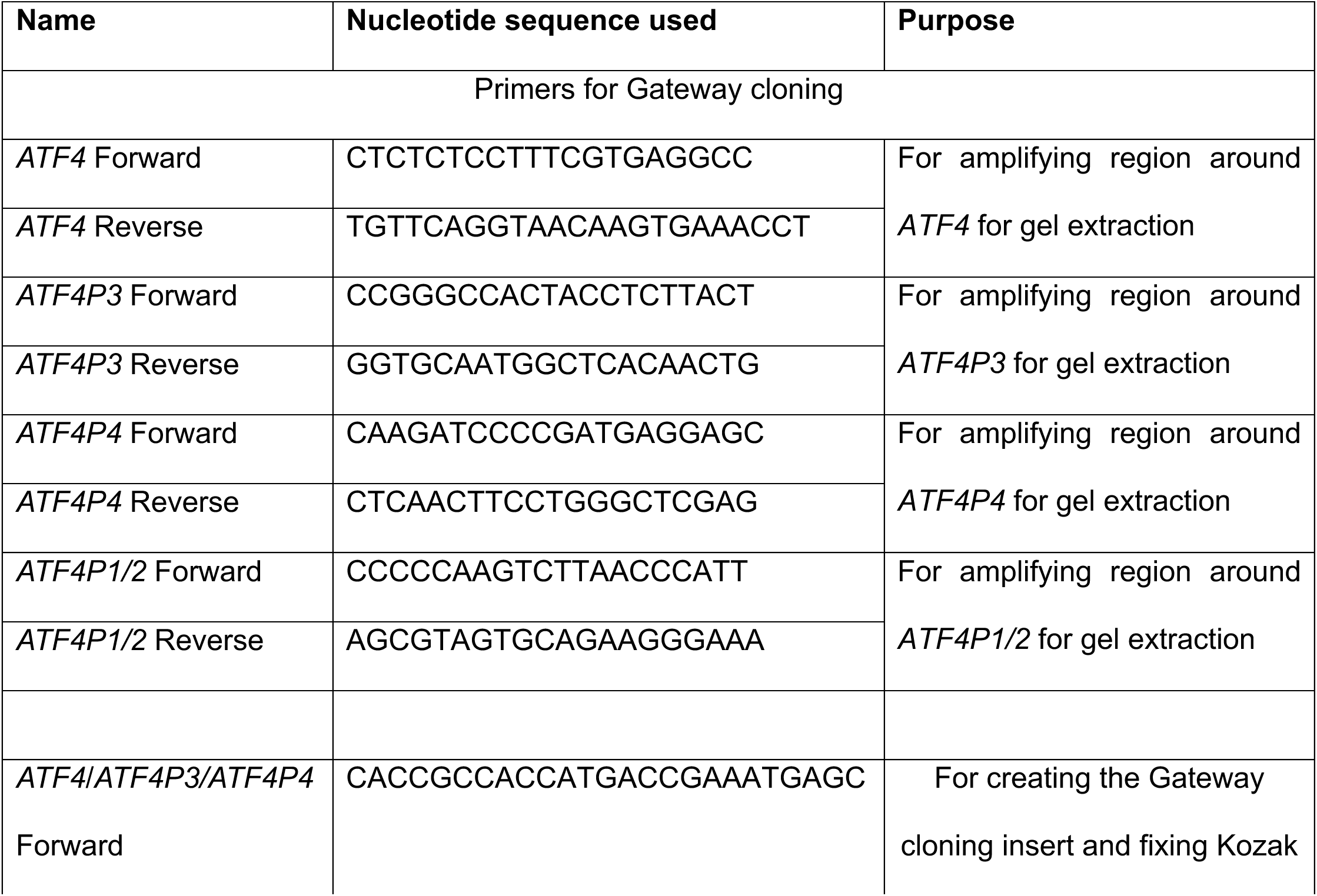
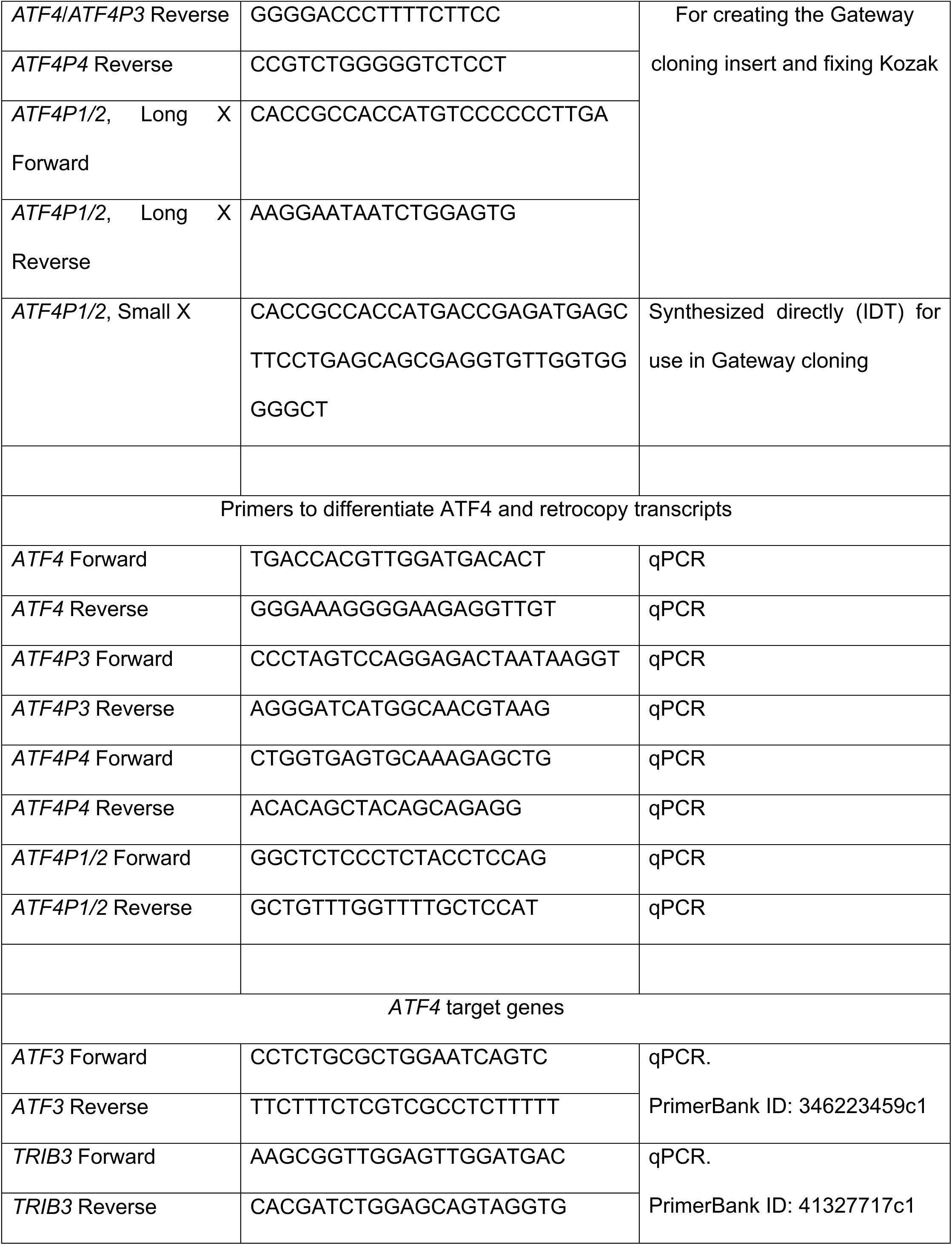
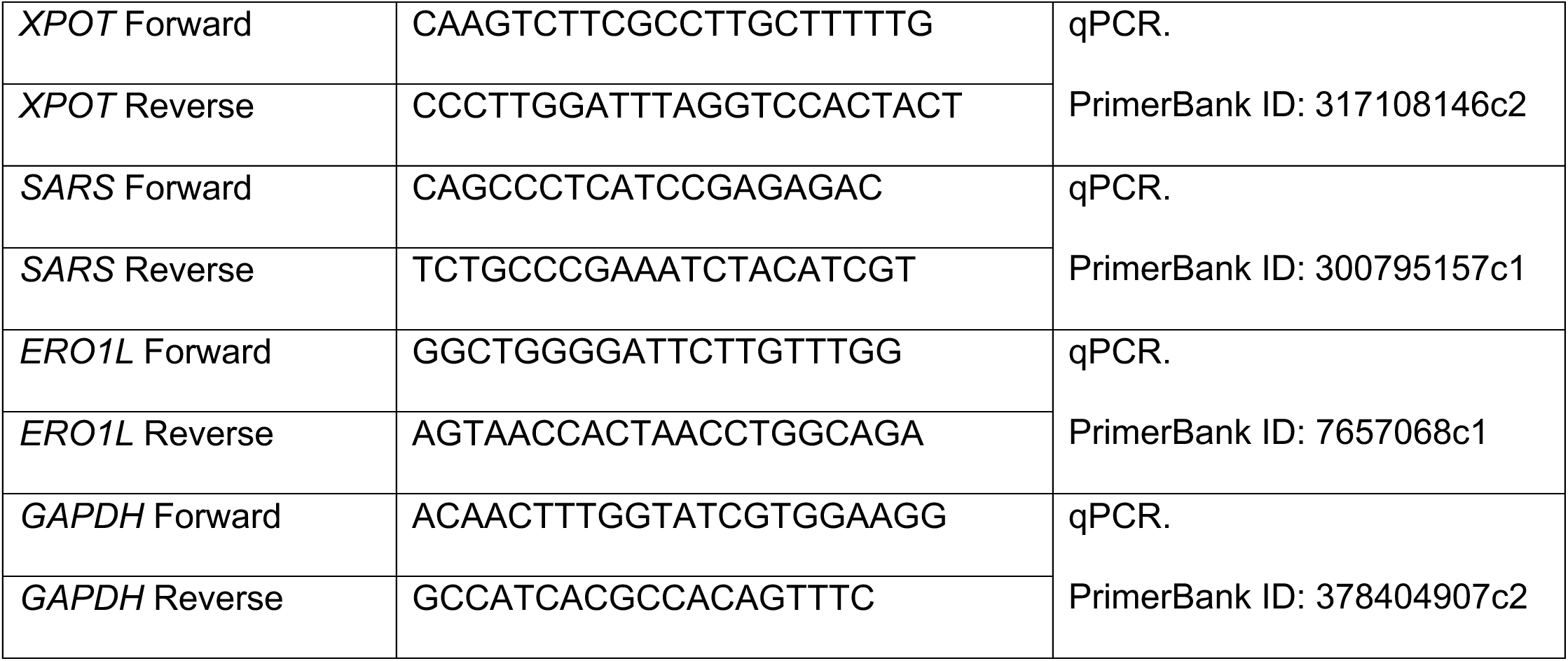

## Supplementary File Legends

**Supp Fig 1.** Mock-up of the genomic region containing the gene *RNF157*. *ATF4P3* was inserted between exons 1 and 2. Image was created using MacVector™ v.18.2.5.

**Supp Fig 2.** Species tree of all organisms that we queried for *ATF4* retrocopies. Various colored shapes represent the status of any *ATF4* retrocopy and/or duplication found upon searching BLAT (with a BLAT score of at least 100). *Catarrhini* retrocopies, containing all retrocopies of *ATF4* found in humans, are contained in a blue box. In this box, checkmarks represent the presence of an *ATF4* retrocopy being found syntenic to *ATF4P1-P4*, or independent *ATF4* retrocopies that are neither syntenic nor duplications of *ATF4P1-P4* (letters A-F). Outside of this box, each horizontal box represents independent retrocopies of *ATF4* (not related to *ATF4P1-4*), while the single vertical box (labeled *Bovidae* retrocopy) is another syntenic retrocopy of *ATF4*. For example, for *Carlito syrichta*, it has four independent retrocopies unique to itself (i.e. these are not *ATF4P1-4* and are not syntenic to any other primate *ATF4* retrocopy). We also note two cases where entire genomes were duplicated (and *ATF4* with them) in *Xenopus*^121^ and ray-finned fish^122^. Finally, we note that we searched the listed closest invertebrate homologs in *Drosophila*, *Caenorhabditis*, and *S. cerevisiae*, none of which had retrocopies.

**Supp Fig 3.** Mock-up of ATF4P3 across humans, chimps, bonobos, and gorillas. We compared protein sequences to the species-specific ATF4 parent gene to determine amino acid changes. Amino acid changes that are shared across all four species are labeled in blue.

**Supp Fig 4.** Graphs of computationally modeled mutation rates of *ATF4* retrocopies. Percentages represent each labeled open reading frame window and are not cumulative of open reading frames greater than that window. Dotted lines represent the approximate evolutionary age of each duplication.

**Supp Fig 5.** UCSC genome browser navigated to *ATF4P3* and *ATF4P4* containing tracks for H3K4Me3 methylation marks on chromatin (red arrow). Note that >2kb on either side display no H3K4Me3 marks, suggesting specificity to each retrocopy.

**Supp Fig 6.** Agarose gel showing the effectiveness of the Zymo DirectZol™ kit at removing genomic DNA from our TriZol™-derived RNA samples of RPE-1 human cells. We used the *ATF4* Forward and Reverse primers listed for qPCR. Lane 1 is a 100 bp ladder (NEB, cat. N3231L) (the two red splotches represent hitting the light threshold for two sizes that are unrelated to the results). Lanes 2-4 are three biological replicates using the DirectZol kit with the optional DNAseI treatment. Lanes 5-7 are three biological replicates the same procedure but omitting the reverse transcriptase reagent. Lanes 8-9 are biological replicates using the DirectZol kit without the DNAseI treatment. If our prepped RNA sample contained residual genomic DNA, that the DNAseI treatment did not destroy, we would have seen bands in lanes 5-7. Thus, we conclude that the DNAseI treatment does fully remove residual genomic DNA, and that lanes 2-4 (and the data in Fig. 3) are derived from actual expressed mRNA. Lanes 8-9 are for reference only to show that DNAse I does have an effect on removing DNA.

**Supp Fig 7.** Images of raw Western blots. A. Western blot of protein extracts from each OE ATF4 cell line using the anti-V5 antibody with no treatment(s). B. Uncropped Western blot of Figure 4. We note here that Btz treatment did not upregulate a visible band for the ATF4P1/2 Small X OE cell line. C. A second biological replicate of a Btz-treatment Western blot, as in (B) and Figure 4. This blot was globally contrast-adjusted to visualize fainter bands. D. Sequenced plasmid insert of ATF4P4 overexpression clone, starting at the ATF4P4 ORF. Note that the ladders are part of the original blot (and are overlaid in their general position on the blot) but are imaged using a different fluorophore, hence the difference in contrast in the ladder.

**Supp Table 1.** Table listing all retrocopies (and their duplications) found in *Catarrhini*, included labeled start/stop sites and target-site duplications. Each tab is a different gene, with *ATF4* also listed for reference.

**Supp Table 2.** Document containing estimations of ancestral versions of *ATF4* derived from Ape transcripts to represent the *ATF4P3/4* ancestral gene (14.8 MYA) or *Catarrhini* transcripts to represent the *ATF4P1/2* ancestral gene (37.5 MYA).

**Supp Table 3.** Table listing all observed retrocopies of ATF4 among 97 different species. Each annotated retrocopy had to reach a BLAT score of at least 100 and contain no introns and/or a PolyA tail to be considered a “yes”. Alternatively, we assigned “possible” when one of these features was present, but the sequence was either highly degraded or missing (such as in an incomplete scaffold assembly). Sequences containing many hits from BLAT of a repeat sequence (likely from a transposon) are marked as “Multiple complex duplications”. Non-retrocopy gene duplications are also noted in under “comments”.

**Supp Table 4.** Table of median transcripts-per-million (TPM) of each gene taken from GTEx (www.gtexportal.org). Tissues reaching expression of at least 0.5TPM are highlighted in green, and these data are quantified in a table below. The right table sorts each tissue by the highest expression of *ATF4* to compare retrocopy expression rates.

**Supp Table 5.** Raw Ct values for qPCR analyses of RPE-1 cells under thapsigargin (Tg) or arsenite (As) treatment. Highlighted cells indicate a single cell population with a stronger overall response (in every gene tested) relative to the other populations. “No template” control results are on the right, indicating the maximum possible Cts achieved with no template input (Undetermined means no signal after 40 cycles).

## Acknowledgements

We thank Erik Eide and Dr. Charles Murtaugh for their help in lentiviral cloning, Western blots, and use of equipment. We thank Dr. Nels Elde and Diane Downhour for their samples and guidance in retrocopy experiments and strategies. We thank the HSC DNA Sequencing Core (director Dr. Derek Warner) at the University of Utah for their help in sequencing analysis. We thank Mr. Ben Anderson for his help in manually collecting GTEx data.

