## Supplementary figures and images for "The stress-induced transcription factor ATF4 has multiple conserved retrocopies that can alter gene expression"

### Supp_Fig_1

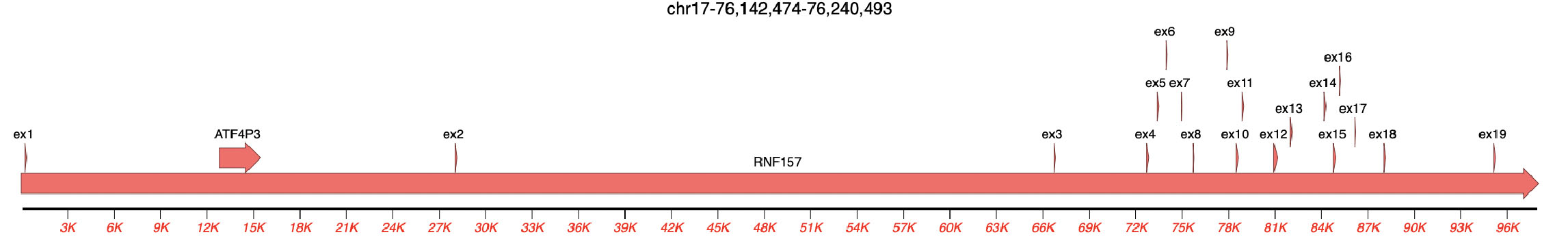

### Supp_Fig_2

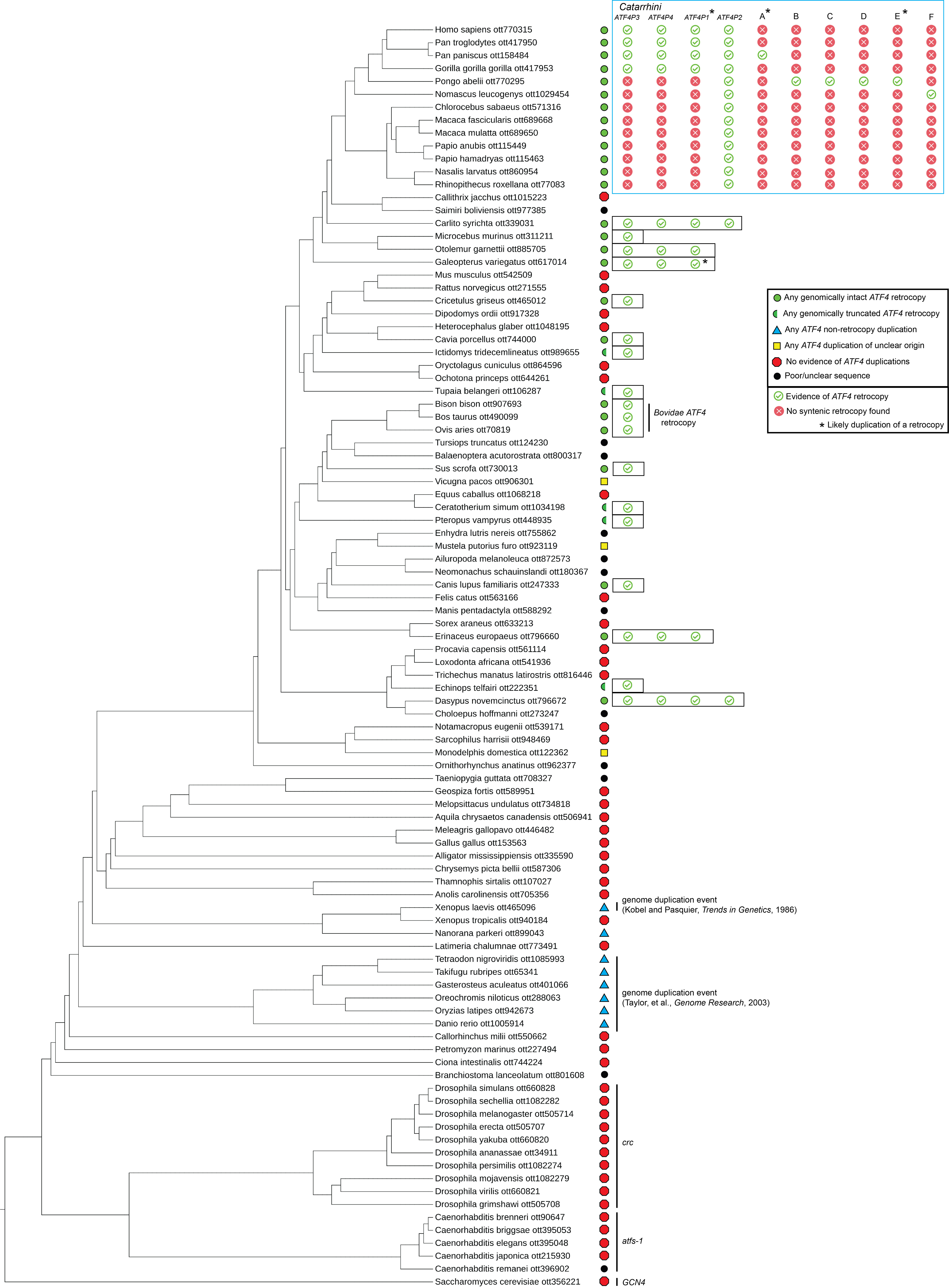

### Supp_Fig_3

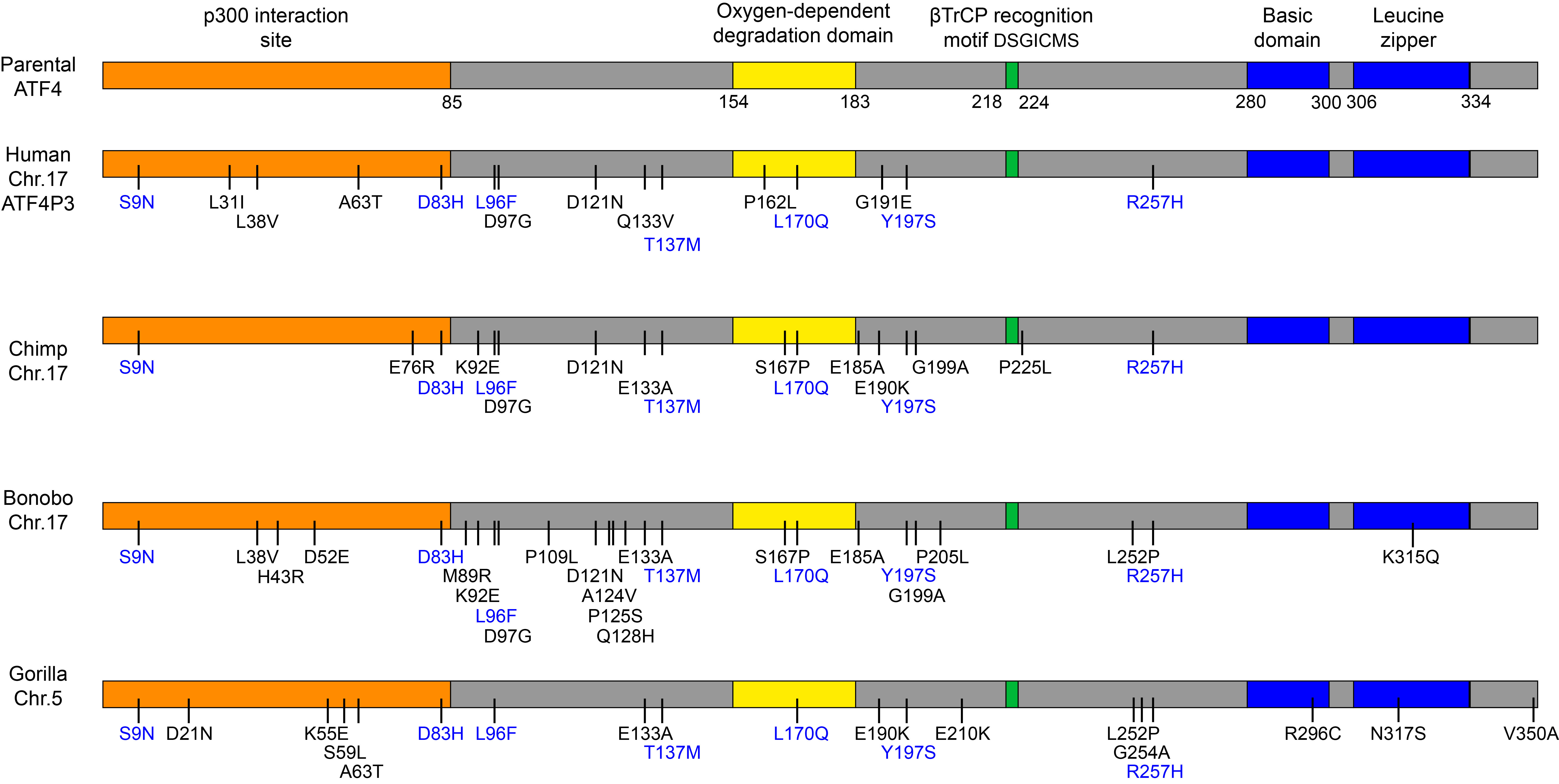

### Supp_Fig_4

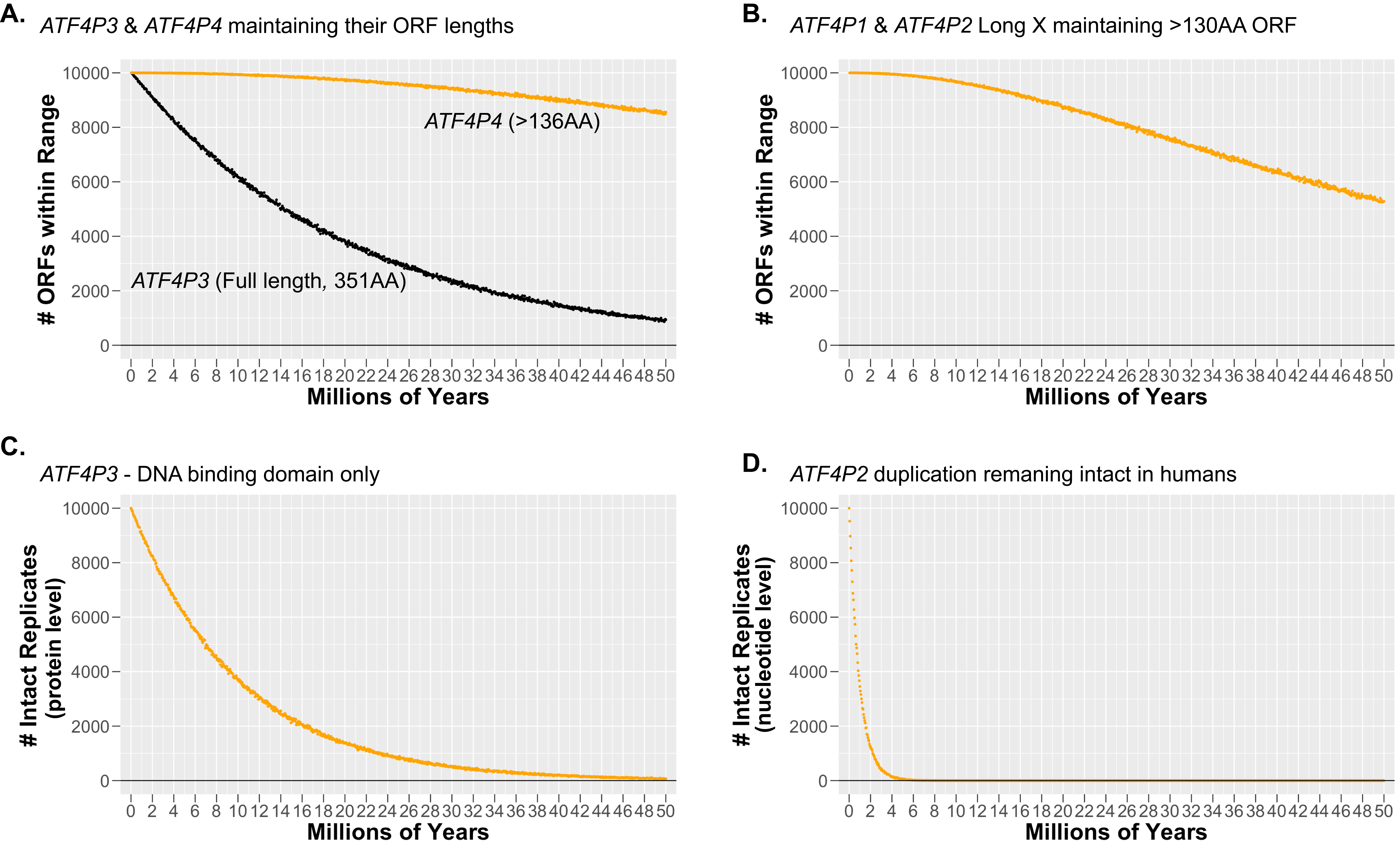

### Supp_Fig_5

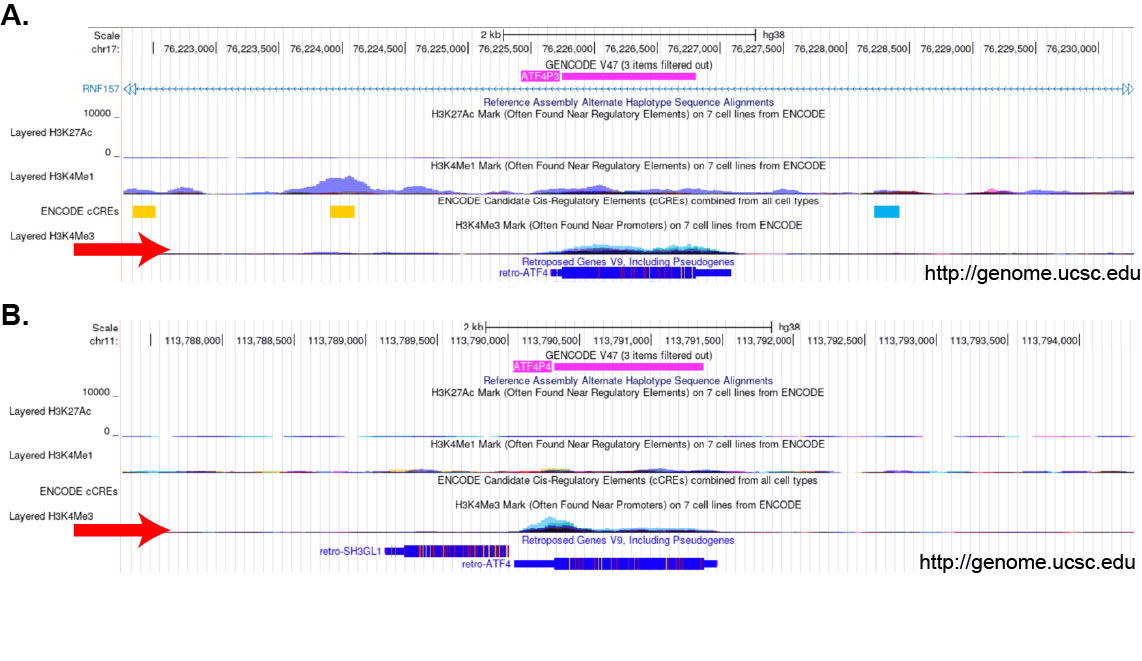

### Supp_Fig_6

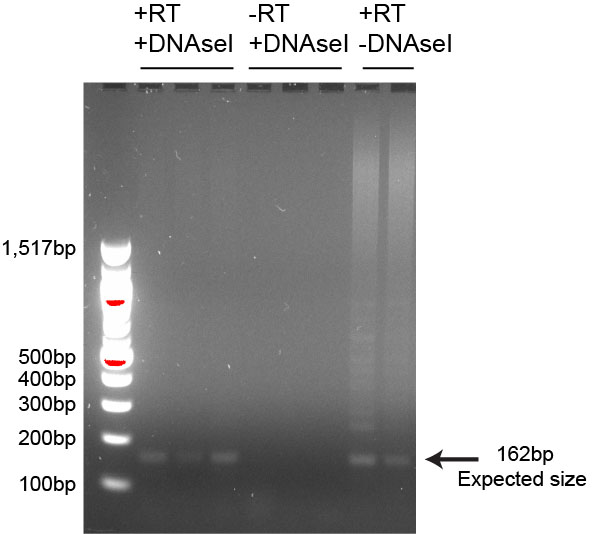

### Supp_Fig_7

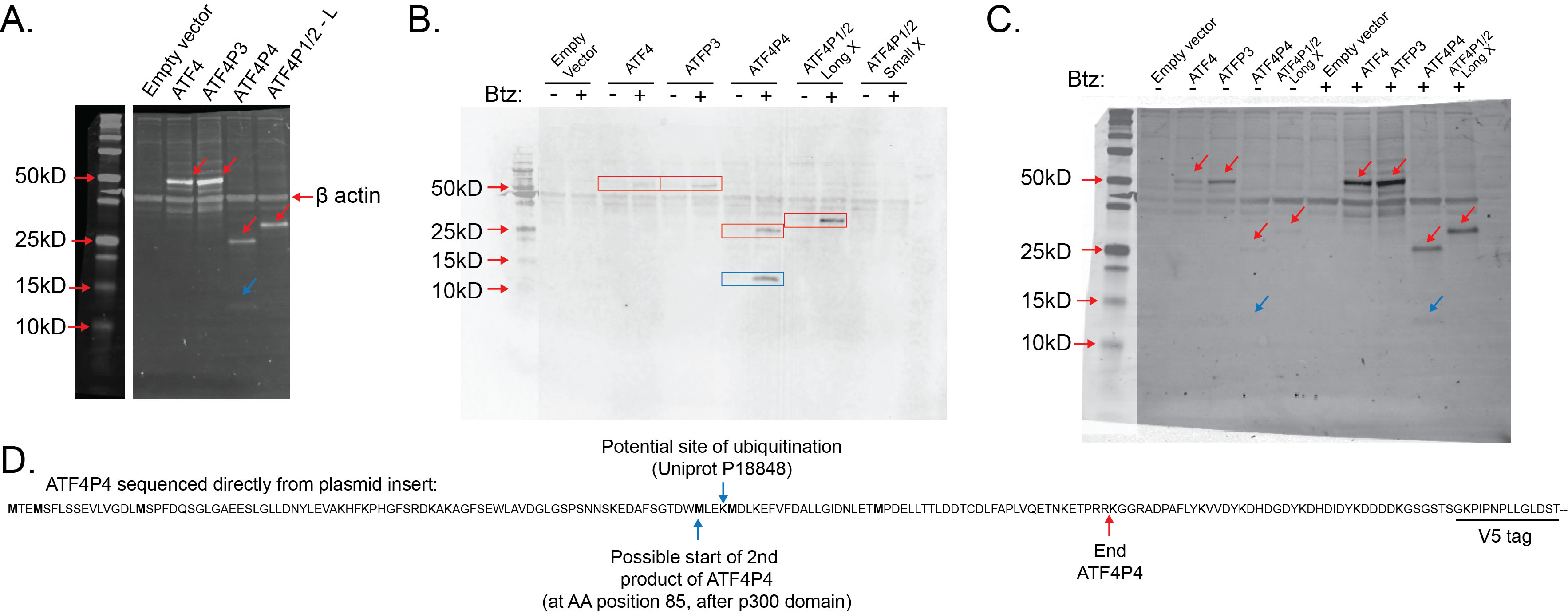
